# pH controlled histone acetylation amplifies melanocyte differentiation program downstream of MITF

**DOI:** 10.1101/545392

**Authors:** Desingu Ayyappa Raja, Vishvabandhu Gotherwal, Yogaspoorthi J Subramaniam, Farina Sultan, Archana Vats, Archana Singh, Sridhar Sivasubbu, Rajesh S Gokhale, Vivek T Natarajan

**Affiliations:** CSIR-Institute of Genomics and Integrative Biology, Mathura Road New Delhi - 110020; Academy of Scientific and Innovative Research, CSIR Road, Taramani, Chennai – 600 113; National Institute of Immunology, Aruna Asaf Ali Marg, New Delhi - 110067

**Keywords:** pH regulation, melanogenesis, p300/CBP, histone acetylation, pigmentation, cell differentiation, epigenetics and zebrafish

## Abstract

Tanning response and melanocyte differentiation are mediated by the central transcription factor MITF. Enigmatically, these involve rapid and selective induction of melanocyte maturation genes, while concomitantly maintaining the expression of other effectors. In this study using cell-based and zebrafish model systems, we elucidate a pH mediated feed-forward mechanism of epigenetic regulation that enables selective amplification of melanocyte maturation program. We demonstrate that MITF activation directly elevates the expression of Carbonic Anhydrase 14 (Ca14) enzyme. Nuclear localized Ca14 increases the intracellular pH, resulting in the activation of histone acetyl transferase activity of p300/CBP. In turn enhanced H3K27 histone acetylation marks of select differentiation genes facilitates their amplified expression by MITF. CRISPR-mediated targeted missense mutation of CA14 in zebrafish results in immature acidic melanocytes with decreased pigmentation, establishing the centrality of this mechanism in rapidly activating melanocyte differentiation. Thereby we reveal a novel epigenetic control through pH modulation that reinforces a deterministic cell fate by altering chromatin dynamics.

## Introduction

Gene expression networks as well as upstream pathways that govern their expression are well-studied for skin melanocytes (Bennett, 1983). Precursors of these cells are neural crest derived and are present in several locations in skin such as hair follicles, dermis and presumably the epidermis (Mort et al, 2015). In response to specific cues from the wnt and melanocortin pathways, these precursors are believed to migrate and rapidly mature into pigmented melanocytes in skin (Harris & Erickson, 2007). Microphthalmia associated transcription factor (MITF), the central melanocyte specific transcription factor is crucial for almost all aspects of development, maintenance and survival of melanocytes across vertebrates (Levy et al, 2006).

To mediate these wide ranging effector functions, the MITF target genes are selectively activated during specific melanocyte transitions (Levy & Fisher, 2011). The level and activity of MITF is governed by cAMP/PKA and MAPK signaling pathways respectively (Johannessen et al, 2013). M/E-box mediated direct activation is observed in several of the MITF target genes. Physical interaction of MITF with factors such as YY1 and proximal interplay with Sox10 mediates activation of a subset of these effector genes, thereby adding another layer of complexity to the selective activation (Li et al, 2012). Recruitment of transcriptional co-regulators such as BRG1 and p300/CBP facilitates dynamic interaction of MITF to subset of promoters (Sato et al, 1997). A key challenge still is to selectively activate certain gene modules dynamically, while maintaining other effector functions of MITF (Li et al, 2012; Praetorius et al, 2013). This is evident during tanning response and melanocyte differentiation wherein MITF activates pigmentation genes several folds, while maintaining the expression of proliferation and survival genes (Malcov-Brog et al, 2018). These recent findings highlights that multiple linked regulatory loops enable selective outcomes in pigmentation during tanning response. Basis of this selectivity by MITF and the mechanistic understanding is just beginning to emerge, wherein epigenetic factors are thought to play a class-specific activator role (de la Serna et al, 2006; Keenen et al, 2010; Laurette et al, 2015; Malcov-Brog et al, 2018). In this context, experiments by Laurette et al, followed by analysis by Malcov-Brog et al recently identified hightened H3K27 acetylation pattern in the pigmentation genes downstream of MITF.

Epigenetic regulation is likely to be intricately linked to cellular cues that cooperate with external signals in modulating networks involved in determining cell fates. pH homeostasis is a crucial biological process that is linked to several of the cellular pathways and is a plausible candidate (Boron, 2004) (Simons et al, 2009; Tatapudy et al, 2017). While the centrality of pH balance in cell homeostasis is well appreciated, emerging evidence indicate that pH could signal cellular events by programmatically modulating existing networks (McBrian et al, 2012; Tatapudy et al, 2017). An increase in the intracellular pH is critical for the differentiation of mouse embryonic stem cells as well as drosophila adult follicle stem cells, highlighting the role of pH in controlling key cell fates (Ulmschneider et al, 2016).

A large family of carbon dioxide metabolizing enzymes, carbonic anhydrases (CA) regulate pH across life forms (Lindskog, 1997). Members of this family are ubiquitously expressed in many cell types and localize to distinct subcellular compartments (Karler & Woodbury, 1960; Reibring et al, 2014). CA14 is a type I transmembrane protein expressed in a variety of cell types but has been studied in retina, brain, kidney, smooth muscle and cardiomyocytes (Fujikawa-Adachi et al, 1999; Kaunisto et al, 2002). Studies using knock-out mice have established a role of CA14 in buffering alkaline shifts in brain (Shah et al, 2005). Intracellular expression of CA14 in the sarcolemma of smooth muscle cells is known to modulate impulse induced muscle contractions (Wetzel et al, 2007). Cells of the Retinal Pigmented Epithelium (RPE) demonstrate high expression of CA14 on the apical region and *Ca14* knock out mouse is deficient in eliciting a functional retinal light response (Ogilvie et al, 2007).

The emerging link between pH and melanin synthesis is several fold. The enzymes involved in melanogenesis namely tyrosinase (Tyr), tyrosinase related protein (Trp1) and dopachrome tautomerase (Dct) that reside within the melanosmes is modulated by the luminal pH. V-type ATPases are involved in maintaining an acidic pH of these lysosome related organelles. Optimal pH for melanogenesis has remained controversial and marginal luminal alkalinization by protonophores is thought to promote melanin synthesis (Watabe et al, 2004). pH changes in the endolysosomal compartments also alter the trafficking and maturation of melanosomal enzymes in addition to their activity, as well as the type of melanin being synthesized (Halaban et al, 2002), (Bellono et al, 2014), (Wakamatsu et al, 2017). Predictable alterations of the cellular pH modulate the resultant pigmentation in melanocytes. The effect of which is surprisingly high with a robust alteration in the net melanin synthesis. Therefore we decided to systematically study the interplay of pH on the entire melanogenic program at various levels.

In this study we demonstrate that intracellular pH is a critical cue for amplifying the melanocyte maturation program. We trace genes that follow a concordant pattern of regulation with pigmentation and identify Carbonic Anhydrase 14 to be a MITF regulated gene. CA14 acts as a feed forward activator of MITF regulatory network and amplifies the melanocyte maturation gene expression by altering the histone acetylation marks via a programmed intracellular pH change. Using cell-based and zebrafish model systems we demonstrate that the feed forward loop involving Ca14 is critical to mediate the melanocyte maturation program downstream of MITF.

## Results

### Alkaline intracellular pH induces pigmentation by enhancing the melanogenesis gene expression

During the culture of B16 cells in DMEM medium CO_2_-HCO_3_^-^ buffering system maintains media pH. Hence we resorted to establish the pH-pigmentation link by modulating the prevailing CO_2_ levels. We set up progressive pigmentation using the B16 cells as described earlier (Natarajan et al, 2014). In this set up the cells progressively activate the melanogenesis gene expression program and increase pigmentation over a course of 8 to 12 days. To alter the pH, 10% CO_2_ levels were tested. We measured the extracellular pH (pH_e_) using the standard pH electrode under controlled conditions of temperature and CO_2_ saturation. While the cells grown in 5% CO_2_ showed an increase in pH_e_ from around 7.4 to 7.8 on day 4 after induction of the pigmentation program, the 10% CO_2_ sustained a constant pH of around 7.4 **(Supplementary Fig 1A)**. This trend was reflected in the intracellular pH (pH_i_), which is close to 7.9 on day 4 under the routine 5% CO_2_ condition. However, under the 10% CO_2_ the pH remained close to 7.0, similar to the day 0 where the cells are depigmented (**Fig 1A**). When the cells were assessed for the cumulative accumulation of melanin on day 8, under the 10% CO_2_ condition we observed depigmented cells (**Fig 1B**). Melanin content assay performed using synthetic melanin standard confirmed that the level of melanin is significantly low **(Supplementary Fig S1B)**. Further electron microscopic evaluation of day 8 cells confirmed that indeed melanin laden stage III and IV melanosomes are dramatically reduced under these conditions (**Fig 1C**).

**Fig 1:**
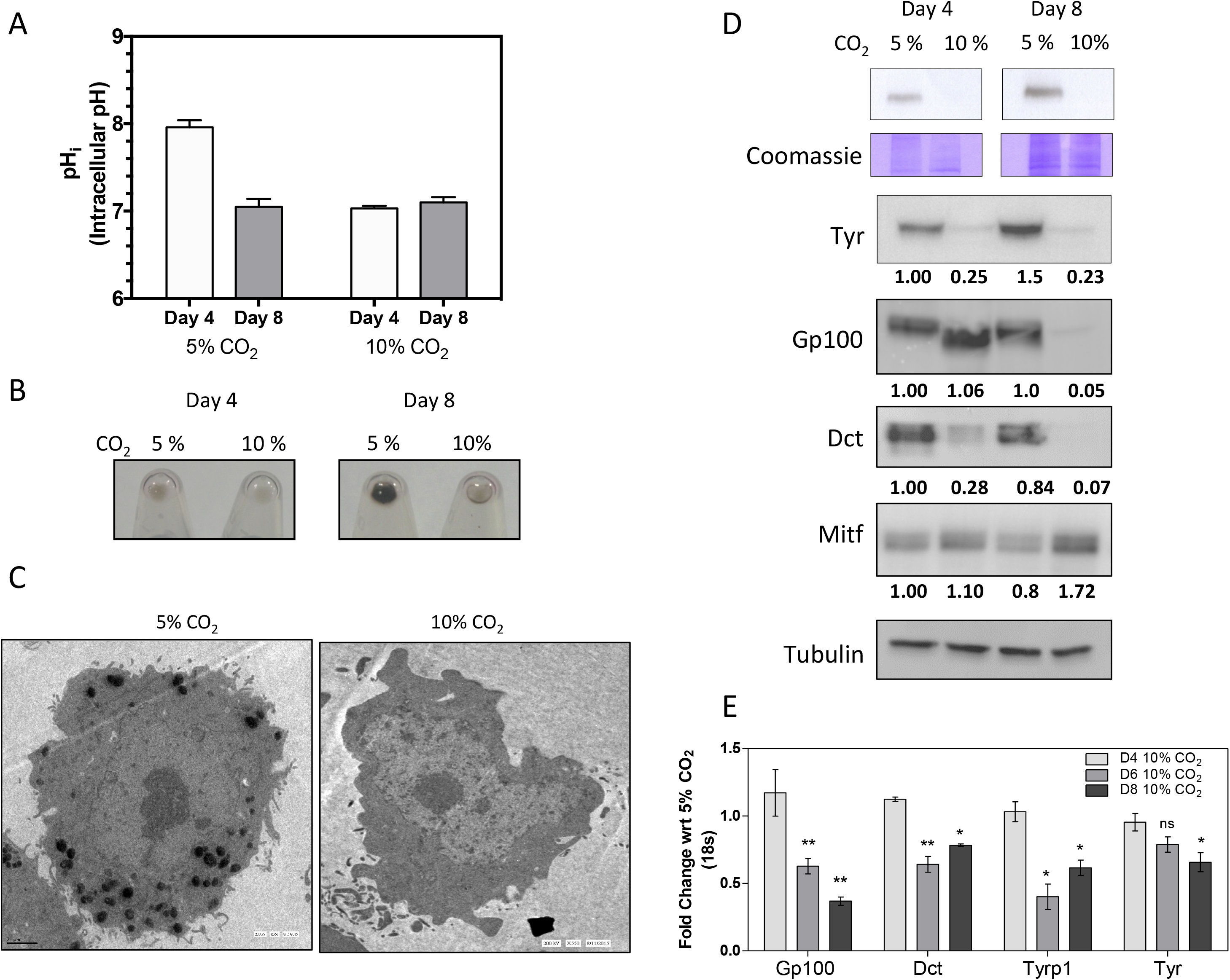
Modulation of intracellular pH by CO_2_ results in predictable changes in pigmentation via a transcriptional change. A.B16 cells were maintained at 5 or 10% CO_2_ for indicated days and intracellular pH (pH_i_) was measured by ratiometric imaging using BCECF-AM for the respective conditions. While the day 4 and day 8 of 10% CO_2_ grown cells remained close to 7.0, in 5% condition the pH_i_ on day 4 was high by almost one pH unit. Data is obtained from three biological replicates with around 60 cells each. B.Pellets of B16 cells at various days under 5 or 10% CO_2_ culture conditions. While the initial day 4 pellets look comparable, but on day 8, 5% CO_2_ grown cells accumulate melanin, while the 10% CO_2_ grown cells remain depigmented. C.Transmission electron micrograph images of day 8 cells indicate that the 5% CO_2_ grown cells have many darkly stained melanosomes, whereas the 10% CO_2_ grown cells are devoid of these pigmented structures. D.Top panel shows *in-gel* tyrosinase activity developed using L-DOPA as the substrate and below is part of the gel stained using coomassie brilliant blue. (bottom panel) Western blot analysis of Tyr, Gp100, Dct, Mitf and Ca14 proteins normalized to tubulin. Numbers represent tubulin normalized fold changes corresponding to day 4 cells grown at 5% CO_2_. 10% CO_2_ reduces expression of pigmentation related markers without a significant decrease in Mitf. E.qRT-PCR analysis of pigmentation related gene transcripts Gp100, Dct, Tyrp1 and Tyr normalized to 18s rRNA on days 4, 6 and 8 during pigmentation. The fold changes are depicted for the corresponding days for cells grown at 5% CO_2_. Progressive reduction in the mRNA expression of pigmentation genes is observed at the transcript level.

We then analyzed steady state protein levels of the components of melanosomal machinery. All the three pigmentation proteins Tyr, Dct and Gp100 were drastically reduced under 10% CO_2_, however the levels of MITF were not decreased, rather we observed a mild increase in the level of this central transcription factor (**Fig 1D**). In concordance with earlier studies that suggested a pH dependent alteration in the protein stability, the activity of tyrosinase enzyme assessed by L-DOPA based in-gel assay as well as western blot analysis confirmed a dramatic reduction (Halaban et al, 2002). However a decrease in Dct and Gp100 levels was unanticipated. Gp100 showed a greater decrease on day 8 and had comparable level on day 4 with alterations in the mobility suggesting processing differences (Hoashi et al).

Surprisingly, transcript levels of these downstream pigmentation genes were lower in 10% CO_2_ condition (**Fig 1E)**. We therefore identified that retaining the cells in an acidic state of pH_i_, suppresses pigmentation by the decreased expression of pigmentation genes despite comparable levels of MITF. We also observed an increase in the cell numbers at 10% CO_2_ **(Supplementary Fig S1C)**. Therefore there is seemingly an additional level of cellular program by intracellular pH that governs melanogenesis beyond enzyme activity and protein stability, involving transcriptional regulation.

### A candidate effector CA14 follows a concordant expression pattern with pigmentation genes

We set out to identify the molecular mechanism behind the pH dependent transcriptional response. Based on our earlier work, we had established the B16 cell autonomous model and demonstrated the utility of this model to identify underlying networks that govern pigmentation. The other model involves the growth of pigmented melanoma tumor derived from mice and grown *in vitro* for four consecutive passages. A set of genes showed a concordant regulation across the two reversible models (**Supplementary Fig S2**). Among the common set of regulated genes, five of the fifty candidates, *Tyr, Tyrp1, Mlana, Mcoln3, Si* and *Rab27a* are targets of the central melanocyte transcription factor MITF. These are well-known pigmentation genes directly involved in the process of melanogenesis and melanosome maturation, which are directly linked to the process of melanocyte differentiation. From this analysis Carbonic anhydrase 14 emerged as a putative candidate gene that could directly regulate pH, as it is regulated with pigmentation in both the cellular model systems.

We then analyzed the pattern of expression of carbonic anhydrases from the data and observed that of the several CAs expressed in melanocytes, only CA14 showed regulation pattern similar to the melanocyte differentiation genes *tyr, dct* and *tyrp1* based on microarray studies (**Fig 2A**) (Natarajan et al, 2014). Hence we proceeded to characterize the regulation of Ca14 with an aim to establish its role in melanocyte maturation.

**Fig 2:**
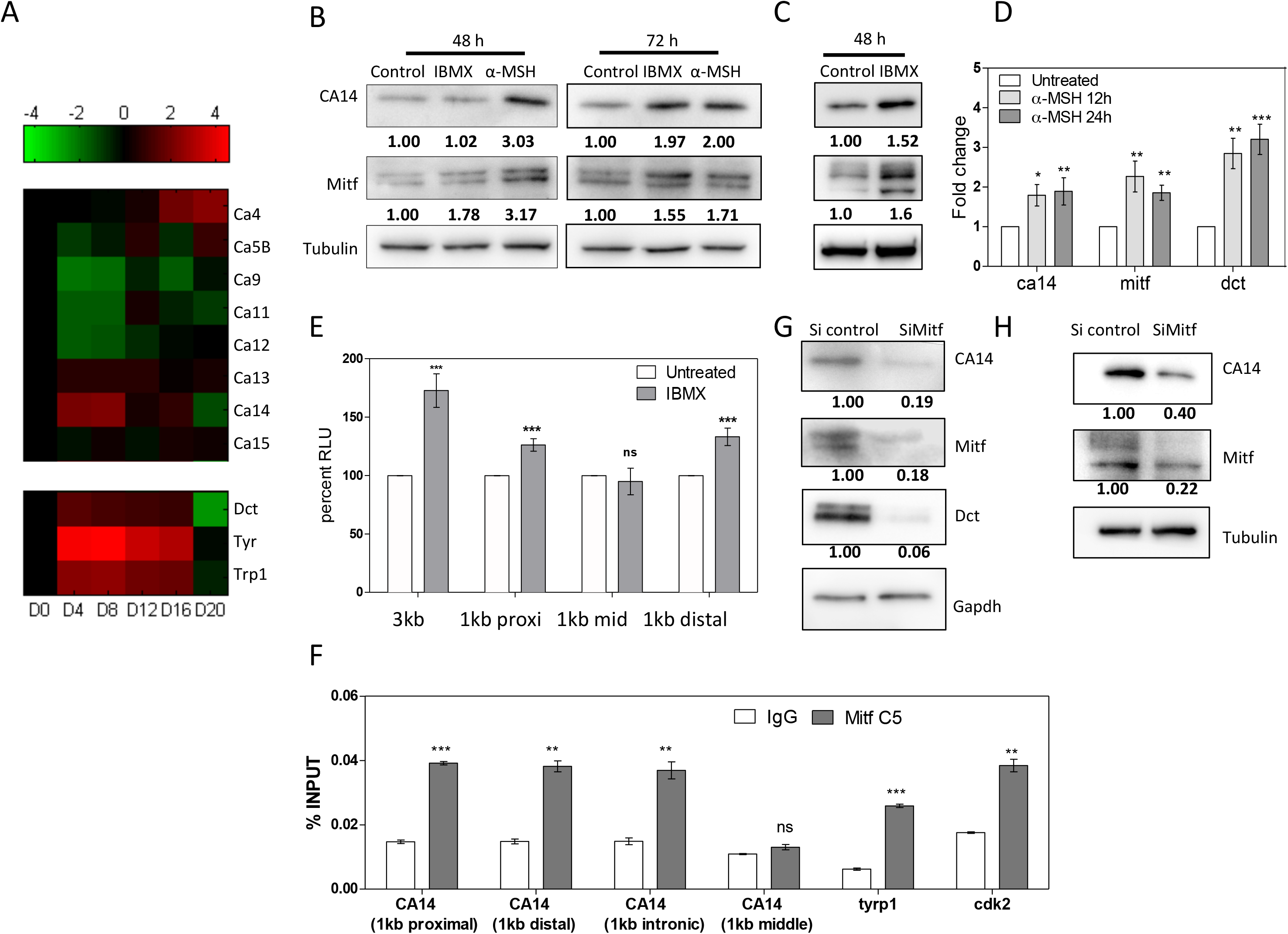
CA14 expression is directly controlled by Mitf. A. (top) Panel of carbonic anhydrases expressed in B16 cells along with (bottom) pigmentation genes during the course of in vitro pigmentation oscillation. The expression changes are represented as a heat-map relative to the expression on day 0. Notably Ca14 follows a concordant pattern of expression with the pigmentation genes. B.Western blot analysis of CA14, Mitf and normalization by Tubulin, on treatment with Mitf inducers 60µM Isobutyl methyl xanthine (IBMX) and 600nM alpha-melanocyte stimulating hormone (α-MSH) for 48 and 72h in Melan-A cells. Fold change with respect to control untreated cells normalized to tubulin expression is depicted as numbers below each lane. C.Western blot analysis of CA14, Mitf and normalization by Tubulin, on treatment with MITF inducer Isobutyl methyl xanthine (IBMX) for 48h in primary human melanocytes. Fold change with respect to control untreated cells normalized to tubulin expression is depicted below each lane. D.qRT-PCR for c*a14, mitf, dct, trp1* and *tyr* transcripts upon treatment with α-MSH for 12 and 24h in Melan-A cells. Fold change is depicted (mean ± SEM, n=2) calculated using *18s rrna* as the reference. E.Dual luciferase assay performed with IBMX for 24h on B16 cells transfected with luciferase construct (pGL4.23) containing 3kb upstream region of *ca14* promoter, or the transcription start sites proximal 1kb, middle, or the distal 1 kb region of the promoter. Renilla luciferase driven by cytomegalovirus promoter (pGL4.75) was used for reference. Error bars represent SEM across three independent experiments. IBMX responsive region appears to be in the 1kb proximal as well as distal regions of the promoter. F.Chromatin Immunoprecipitation using Mitf antibody (C5) or normal mouse IgG control, was followed by qRT-PCR performed for mitf binding sites in Ca14 promoter and intron region. Graphs are plotted as percent input. Trp1 and cdk2 were taken as positive control. Bars represent mean ± SEM across two biological replicate experiments. G.Western blot analysis of Ca14, Mitf and Dct normalized to Gapdh upon knockdown using *Mitf* siRNA in Melan-A cells. H.Western blot analysis of CA14 and MITF normalized to tubulin upon knockdown using *MITF* siRNA in primary human melanocytes.

### CA14 expression is induced upon activation of the melanocortin pathway via MITF

As Ca14 showed concordant expression with pigmentation genes and correlated with the activity of MITF, we set out to identify whether *ca14* is indeed a downstream gene and could play an important role in mediating the effects of MITF in melanocytes. Overexpression of Mitf, followed by global gene expression analysis revealed a set of regulated genes (Hoek et al, 2008). Chromatin immunoprecipitation studies have independently identified several promoters occupied by this transcription factor (Strub et al, 2011). Combined analysis of both these approaches resulted in a set of genes enriched in known targets of Mitf, along with several other putative candidates. In this set of genes Ca14 features and hence emerged as a promising downstream target of MITF.

α-melanocyte stimulating hormone (α-MSH) and Iso-Butyl Methyl Xanthine (IBMX) were used as inducers of Mitf to stimulate melanocytes (Motiani et al, 2018). We carried out these experiments in mouse Melan-A cells, as well as the primary human melanocytes. Treatment of Melan-A cells with 60 µM IBMX or 1 µM α-MSH for 48h and 72h resulted in the induction of Mitf and the downstream target gene Dct, confirming activation of the melanocortin pathway. The antibody used to detect CA14 was validated by western blot analysis wherein we could identify the 25 kDa form of CA14 protein. The intensity of the band was reduced upon silencing with a pool of siRNA against Ca14 and increased with expression of CA14 ORF in an expression vector, confirming specificity of the antibody (**Supplementary Fig S3 A & B**). With IBMX as well as α-MSH treatments we could observe elevation of the 25kDa form of Ca14 protein, indicaing that activation of Mitf mediates Ca14 induction **(Fig 2B)**. We observed a similar induction of CA14 in primary human melanocytes treated with IBMX for 48 h (**Fig 2C**).

We performed qRT-PCR analysis to monitor the mRNA levels in B16 cells treated with α-MSH for 12 and 24h, as the transcript-level changes are expected earlier than protein level changes. We observed a robust elevation in the expression of pigmentation gene transcripts *Tyr, Dct* and *Trp1*. Moreover the elevation in *Mitf* and *Ca14* levels were comparable, modest but significant **(Fig 2D)**.

### MITF directly binds to Ca14 promoter and regulates its expression

To probe the induction of *ca14* by Mitf further, we cloned 3 kb upstream region of *ca14* in a reporter vector and performed dual luciferase assays. Substantial elevation of reporter activity with IBMX and α-MSH was observed confirming that the induction to be a transcriptional response **(Fig 2E)**. Analysis of MITF binding sites in the promoters of Ca14 gene indicated multiple sites in both the human and mouse promoter regions (**Supplementary Fig S4**). To fine-map the responsive site we adopted two strategies, in the first strategy we created three clones of mouse Ca14 promoter carrying the 1kb transcription start site proximal region, the mid or the distal 1 kb region **(Fig 2E)**. Luciferase assays confirmed that the IBMX inducibility was present in the proximal and the distal regions whereas the middle 1 kb promoter sequence was not IBMX inducible.

Further, chromatin immunoprecipitation using C5 monoclonal antibody to MITF demonstrated binding to 3kb as well as the proximal and distal regions was observed. Comparable binding of Mitf to the intronic site in Ca14 gene as well *Tyrp1* and *Cdk2* promoters confirmed direct binding of MITF to Ca14 for its regulation. However the middle region did not show appreciable binding of MITF (**Fig 2F**). We propose that there are multiple binding sites within the CA14 promoter responsible for its direct induction by MITF. Finally, Mitf dependence of *ca14* expression was probed by the downregulation of this transcription factor by siRNA followed by western blot analysis. While the known downstream Mitf target gene *Dct* was dramatically downregulated, both Mitf and Ca14 showed a comparable downregulation of around 80% **(Fig 2G)**. Similar observations were made from primary human melanocytes using a siRNA against human Mitf, confirming the Mitf dependency of CA14 expression **(Fig 2H)**. Hence we establish that melanocyte differentiating melanocortin signaling pathway controls Ca14 expression in a MITF dependent manner in both mouse as well as human cells.

### CA14 is essential for melanocyte maturation in zebrafish

In cultured melanocytes pH is governed by prevailing CO_2_ concentrations which may override intracellular pH programs. Therefore to address the role of CA14 on melanocyte functions we utilized a morpholino (MO) based transient silencing approach using the zebrafish model system. Herein the pigment producing cells termed melanophores are ontologically equivalent to higher vertebrate melanocytes and the underlying gene networks are highly conserved. During embryogenesis melanophores rapidly mature between 48 to 72 hours post fertilization (hpf). Additionally in the absence of melanosome transport, melanocyte maturation is easy to visualize and monitor.

After titrating the dose of the Ca14 MO based on viability of embryos, scoring of the phenotypes was carried out **(Supplementary Fig S7)**. Analysis of melanophores at 48 hpf indicated that the *ca14* morphants were lightly pigmented compared to control non-targeting MO-injected embryos at the same concentration (**Fig 3A & C**). To assess melanophore numbers, we generated morphants in transgenic zebrafish line *Tg:ftyrp1-GFP* wherein the melanophores are fluorescently marked with GFP driven by a fugu Tyrp-1 promoter (Zou et al, 2006). To prevent masking of GFP fluorescence by melanin, the embryos were treated with phenylthiourea (PTU), a potent tyrosinase inhibitor. Similar pattern of melanophore positioning and comparable number of melanophores were observed in the morphant fishes suggesting that the melanophore numbers are unaltered which indicates melanocyte specification and survival are unaffected (**Fig 3B** right panel & **3D**). However, the mature heavily pigmented melanophores were drastically reduced in the *ca14* morphants. Quantitation of the bright field images using Image J platform suggested that the morphant melanophores had a higher mean grey value, indicating that they were lighter with less melanin content (**Fig 3C**). These observations strongly suggest that CA14 plays an important role in the process of melanogenesis, a crucial event associated with melanocyte maturation. Furthermore, upstream events such as specification, migration and patterning of melanocytes are seemingly unperturbed. We observed a progressive increase in the expression of pigmentation genes *tyr, dct* and *tyrp1b*, concomitant with the known melanocyte maturation process; however in the ca14 morphants that elevation was severely curtailed (**Fig 5E-G**). Therefore CA14 mediated melanocyte maturation by altering the pigmentation gene expression. We then went ahead to elucidate the mechanism of CA14 mediated pigmentation using cultured cells as the model system.

**Fig 3:**
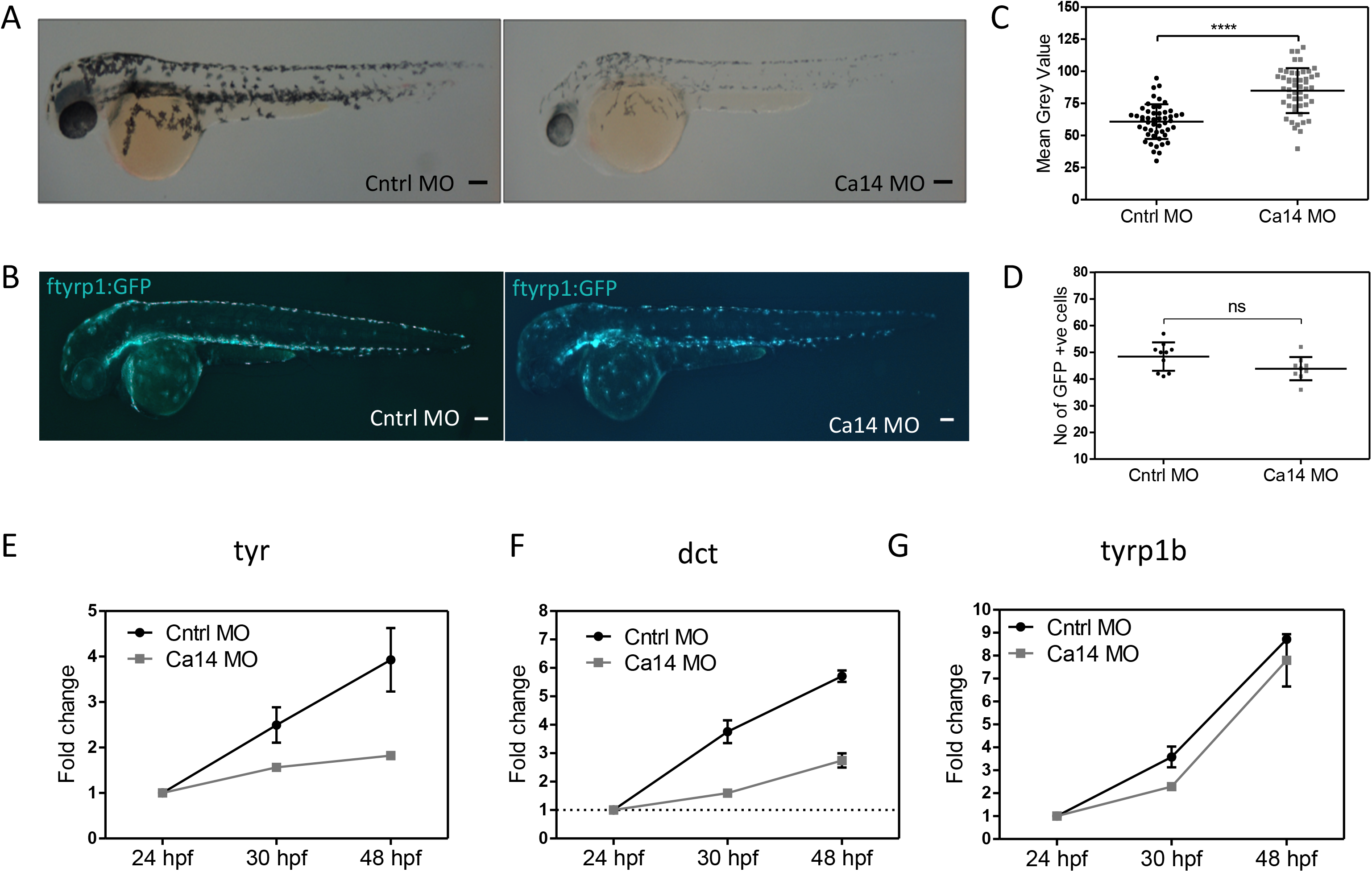
Transient silencing of *Ca14* in Zebrafish decreases melanocyte maturation. A.Brightfield images of the lateral view of control and Ca14 morphant embryos at 48 hours post fertilization (hpf). The black structures observed are melanophores, the melanin containing cells equivalent to mammalian melanocytes. These are pale in the morphants. Scale bar 100 μm. B.Representative fluorescence images of control and Ca14 morphant embryos at 48 hours post fertilization (hpf) from ftyrp1:GFP line, wherein the melanocytes are marked by the expression of GFP. Number of melanocytes remain unaffected in the morphants. C.Melanin quantitation from the bright field images of control and Ca14 morphants carried out using Image J platform. The mean grey values are inversely linked to melanin content of the embryo, and are represented as scatter plot across melanophores from multiple animals. D.Number of ftyrp1 promoter driven GFP positive melanophores from control and Ca14 morphant embryos at 48 hours post fertilization remains unchanged. E.qRT-PCR quantification of pigmentation genes tyrosinase (tyr), F. dopachrome tautomerase (dct) and G. tyrosinase related protein 1b (tyrp1b) between control and ca14 morphants at 36 and 48 hpf, compared to 24 hpf using beta-actin as the normalization reference. Error bars represent SEM across three independent experiments normalized to corresponding control MO set with RPS11 as the normalization control.

**Fig 4:**
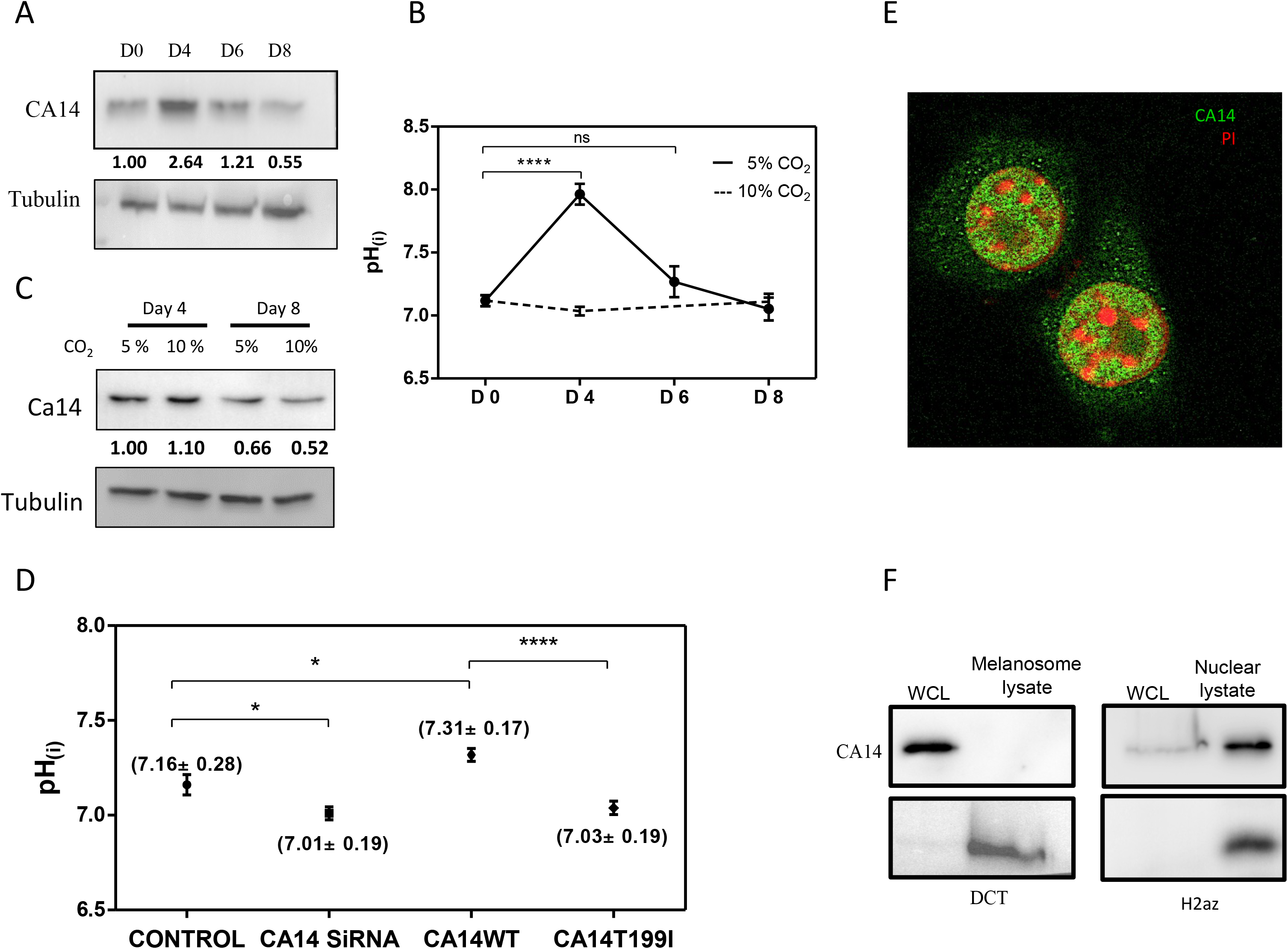
CA14 causes increase in intracellular pH observed during pigmentation program. A.Western blot analysis of CA14 normalized to tubulin during pigmentation. Indicated days (day 0, 4, 6 and 8) represent number of days after initiating the pigmentation program in the pigment oscillator model. Numbers indicate normalized fold change *wrt* to tubulin with day 0 as the reference. B.Intracellular pH (pH_i_) probed by ratiometric pH sensitive fluorescent dye BCECF-AM in B16-melanoma cells during different days of pigmentation. The data represents mean ± standard deviation of at least 100 randomly chosen cells. Statistical analysis was performed using unpaired t test across three independent biological replicates. The trend in pH_i_ follows Ca14 expression in 5% CO_2_ condition. While pigmentation conducive 5% CO_2_ shows an increase in pH_i_, the non-conducive 10% CO_2_ does not show the elevation in pH_i_. C.Western blot analysis of Ca14 normalized to tubulin on day 4 and day 8 of pigmentation from cells grown at 5% and 10% CO_2_ respectively. Please note that the normalization blot for tubulin represented here is also depicted in Fig 1D. The increase in Ca14 expression on day 4 is evident under both conditions, thereby prevailing CO2 levels in this system drives pH_i_ changes. D.pH_i_ probed by BCECF-AM in B16-melanoma cells on siRNA knockdown of *ca14* and on overexpression of wild type or the catalytically inactive form (CA14_T199I_). The data represents mean ± standard deviation of at least 30 transfected cells. Statistical analysis was performed using unpaired t test. The data is representative of two experimental sets. E.Confocal image of immunocytochemistry using CA14 antibody (green) on permeabilized B16 melanoma cells, counter stained by PI (red). Nuclear localization of the Ca14 signal is evident. F.Western blot analysis of subcellular fractions enriched in nucleus and melanosome, using CA14 antibody. H2AZ antibody was used as a marker for nuclear and DCT for melanosomal enrichments.

**Fig 5:**
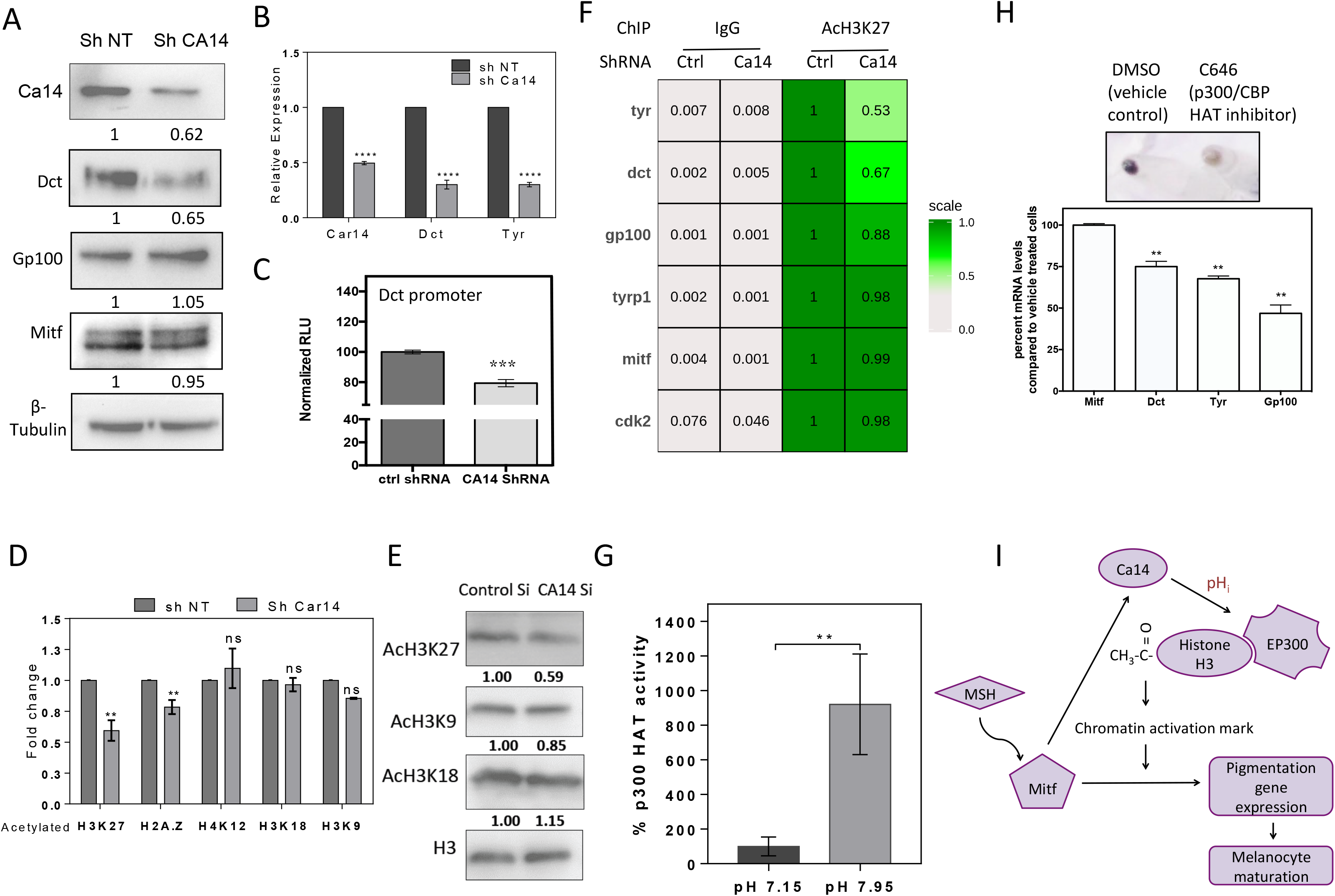
CA14 brings about transcriptional regulation of downstream pigmentation genes through histone acetylation. A.Western blot analysis of Dct, Gp100 and Mitf proteins along with Ca14 and tubulin upon knockdown using *Ca14* shRNA in B16 cells. Numbers represent tubulin normalized fold changes wrt control non-targeting ShRNA. B.Gene expression analysis using qRT-PCR of Ca14 and Dct, upon *ca14* knockdown using shRNA in B16 cells. Fold change normalized to gapdh wrt control non-targeting ShRNA. Bars represent mean ± SEM across 4 independent biological replicates. C.Luciferase assay of *Dct* promoter cloned downstream of firefly luciferase (pGL4.23), on knockdown using *ca14* shRNA. Bars represent mean ± SEM across 3 independent biological replicates. D.Western blot analysis was carried out for specific acetylation of histones (H3K27, H2A.Z, H4K12, H2AK5 and H3K9) in Ca14 knockdown B16 melanoma cells. Bars represent log normalized fold changes wrt control non-targeting shRNA of mean ± SEM across 2 independent biological replicates. E.Representative western blot of acetylations in histone H3. F.Chromatin Immunoprecipitation using acetylated H3K27 antibody or normal rabbit IgG as a control on day 4 pigmenting cells of the oscillator in control non-targetting ShRNA and Ca14 shRNA transfected cells. qRT-PCR was performed for select promoters in the immunoprecipitate and input DNA. Relative normalized percent input DNA under each promoter is depicted as a heat-map. G.Histone acetyl transferase (HAT) activity as a function of pH for recombinant HAT domain of p300 protein is depicted. Compared to the HAT activity at pH 7.15, the activity at pH 7.95 was almost 8 fold. H.Cell pellets and real-time qRT-PCR levels of pigmentation transcripts Dct, Tyr, Gp100 and MITF upon selective inhibition of p300/CBP HAT activity in B16 cells. p300 HAT inhibition resulted in depigmented cells with a decrease in pigmentation gene transcripts. I.Proposed model of melanocyte maturation depicts Mitf mediated induction of Ca14, which increases intracellular pH (pH_i_) and changes chromatin activation marks (AcH3K27) on selective pigmentation genes, amplifying their expression by MITF.

### CA14 increases intracellular pH during pigmentation

Since Ca14 is a MITF dependent gene, expression changes during pigmentation and a downstream modulation of pH due to its carbonic anhydrase activity emerged as a possibility. To study this we induced pigmentation in B16 cells using the pigmentation oscillation. On different days of the pigmentation oscillator the levels of the Ca14 mRNA and protein using q-PCR and western blot analysis respectively was quantified. CA14 expression at both protein as well as mRNA levels was high during early phase (day 4) and progressively decreased at the later days of pigmentation (**Fig 4A**, **Supplementary Fig S5)**.

As carbonic anhydrases mediate pH buffering, we measured pH_i_ using the ratiometric pH sensitive dye (BCECF-AM) that reports intracellular pH (pH_i_). We observed a consistent increase in pH_i_ from around 7.0 on day 0 to 7.9 on day 4, which is subsequently restored to around 7.0 on day 8 of the oscillator (**Fig 4B**). Strikingly, this trend followed changes in Ca14 expression. The pigmentation non-permissive condition involving 10% CO_2_ did not demonstrate a sharp rise in pH_i_ on day 4. We observed a decrease in Ca14 protein levels on day 8 in both 5% and 10% conditions, suggesting that CO_2_ mediates its effects on pH by shifting the equilibrium towards an acidic pH (**Fig 4C**). We then proceeded to unequivocally establish the role of Ca14 in altering pH_i_.

We transfected B16 cells with C-terminal mCherry tagged CA14 to trace the transfected cells for pH measurements. There was a marginal but significant elevation of pH_i,_ which could not be observed for the catalytically inactive mutant form (CA14_T166I_). Silencing *ca14* using a cocktail of four independent siRNAs caused intracellular acidification and reduced pH_i_ **(Fig 4D)**. Together the two data confirm the role of CA14 in elevating intracellular pH. CA14 mediated elevation of pH_i_ in B16 cells raised the question about the localization of CA14 in melanocytes.

### CA14 is localized to the nucleus in melanocytes

CA14 is an alpha-type carbonic anhydrase domain containing type I transmembrane protein, preceded by a short signal sequence (Whittington et al, 2004). Hence the catalytic motif is predicted to function in buffering extracellular pH (pH_e_), alternately its localization to intracellular membranes would modulate organelle pH. Expression of Ca14 is high in adult brain, retinal pigmented epithelium, liver, heart, and skeletal muscle and its localization in various cell-types differ (Hallerdei et al, 2010; Purkerson & Schwartz, 2007; Vargas & Alvarez, 2012; Wetzel et al, 2007). While in smooth muscle cells its expression is detected in sarcoplasmic reticulum, in other cell-types expression in plasma membrane is indicated (Wetzel et al, 2007). Being a MITF inducible gene Ca14 like other targets could be localized to melanosomes, where the pH regulation is known to be critical (Ancans et al, 2001). However its control over pH_i_ indicates its localization to the cytoplasmic or a connected subcellular compartment.

Immunofluorescence studies were carried out on B16 cells to identify the localization of CA14 in melanocytes. Intracellular localization of CA14 could be detected in the nucleus of B16 cells (**Fig 4E**). Fractionation of cells to enrich the nuclear and melanosomal fractions, followed by western blot analysis confirmed CA14 localization to the nucleus and not to melanosomes **(Fig 4F)**. This is strikingly similar to another transmembrane paralog CA9 that localizes to the nucleus and is involved in nucleolar gene expression (Sasso et al, 2015). CA9 has been an important target for pH modulation in several cancers including melanoma (Supuran & Winum, 2015).

While culturing primary human melanocytes in the laboratory we had previously observed that the cells grown under MBM-4 medium compared to M254 medium were more proliferative and had less melanin content. Similar observations were also reported earlier (Kormos et al, 2011). We observed that the expression of CA14 was higher in M254 medium and localization was primarily in the nucleus. Whereas in MBM-4 media the cells had lower expression of ca14 and the immuno-localization was found to be diffuse **(Supplementary Fig S6)**. These observations further add credence to a possible role of CA14 in melanocyte maturation. To understand the mechanism we proceeded to investigate the effect of CA14 silencing on the expression of pigmentation genes in B16 cells.

### Ca14 mediates pigmentation gene expression through a transcriptional response

*Ca14* by virtue of being regulated by Mitf, its control over pH_i_ and its role in sustaining melanin content of zebrafish melanophores makes it a likely candidate to mediate melanocytes maturation. Hence we set out to address the effect of down-regulating *ca14* on the expression of pigmentation genes. We transfected B16 cells with a pool of four siRNAs targeting *ca14* or control non-targeting siRNA pool and the cells were subjected to downstream experiments. Independently, shRNA mediated knockdown was also carried out. Western blot analysis was performed on the cell lysates and a reduction of around 60% was observed for CA14 (**Fig 5A and Supplementary Fig S3A**). Expression of pigmentation genes was found out to be lower upon Ca14 silencing in both the approaches. mRNA levels of *dct and Tyr* was found to be downregulated by q-RT PCR analysis (**Fig 5B**). We further confirmed that the down-regulation is at the transcriptional level by performing luciferase assays using *dct* promoter driven firefly luciferase (**Fig 5C**). The Dct promoter activity was marginally but consistently downregulated by around 30% upon silencing of ca14. We observed an increase in the promoter activity of an unrelated promoter kif1b suggesting that the decrease is unlikely due to possible alterations in luciferase activity due to changes in pH.

Given the nuclear expression of CA14, it is likely that the local pH changes may culminate in specific transcriptional response. Since Dct and the other melanocyte differentiation genes are direct targets for MITF, it is surprising that Ca14 is able to modulate gene expression without affecting MITF levels. It is therefore likely that Ca14 could mediate transcriptional activation by facilitating chromatin alterations, which in turn facilitate MITF mediated gene expression. This possibility also allows for promoter specific alterations rendering selectivity in the downstream gene expression. In an earlier work the authors reported changes in histone acetylation upon intracellular pH change (McBrian et al, 2012), hence we set out to investigate this possibility.

### Ca14 promotes H3K27 acetylation marks on select MITF target genes

Thus far we could establish that Ca14 is present in the nucleus, increases intracellular pH and enhances the expression of pigmentation genes. We went ahead and monitored global changes in histone acetylation using western blot analysis and observed a modest but consistent decrease across AcH3K27 (Acetylated Histone 3 at Lys 27 position) and AcH2A.Z, but not in AcH3K18, AcH3K9 or AcH4K12 **(Fig 5D)**. This provided an exciting possibility of local pH changes culminating in epigenetic marks that would in turn affect pigmentation gene expression profiles downstream of MITF. While the trend of decrease in AcH3K27 was consistent across three independent biological replicate experiments, the effect was marginal around 30 – 40% reduction in global acetylation. We then proceeded to query the specific changes in a battery of promoters of MITF target genes using chromatin immunoprecipitation (ChIP).

We transfected B16 cells with control non-targeting or the CA14 targeting ShRNA construct and allowed cells to initiate pigmentation by setting up pigmentation oscillation. Cells were subjected to crosslinking on day 5, a day after the peak in pH_i_ is observed and chromatin immunoprecipitation was carried out with acetylated H3K27 as well as control IgG. Relative enrichment with respect to the input DNA was quantitated by q-RT PCR using promoter specific primers. The enrichment of AcH3K27 was decreased at the promoters of *Tyr, Dct* and *Gp100* genes (**Fig 5F**). We observed that AcH3K27 occupancy at Tyr promoter was reduced by around 50%, *Dct* promoter by 33% and *Gp100* by a marginal but consistent decrease of around 12%. However the promoters of other MITF targets *Tyrp1*, and *Cdk2* as well as the promoter of *Mitf* itself remained unaltered upon Ca14 silencing (**Fig 5F**). This experiment confirmed that Ca14 brings about the promoter specific changes in activation marks and provides a molecular basis of pigmentation gene expression control mediated by MITF. This data also reinforces earlier observations that the pigmentation gene targets of MITF have unusually high H3K27 acetylation profiles as compared to global average or other targets of MITF (Malcov-Brog et al, 2018).

### Histone Acetyl Transferase activity of p300 is elevated at an alkaline pH

We then set out to identify the pH dependent mechanism of histone acetylation. p300/CBP emerged as a probable candidate as it mediates H3K27 acetylation, additionally physical interaction of MITF with p300/CBP is already established (Sato et al, 1997). In vitro activity of recombinant p300 HAT domain was carried out with a synthetic peptide of histone H3 and acetyl CoA, and the product CoA is measured by coupling with a fluorophore. Activity at around the neutral pH 7.15 was 1/8^th^ of its activity at pH 7.95; thereby suggesting that acetylation by p300 would be clearly high at alkaline pH **(Fig 5G)**. Hence this study provided a possible effector of the pH_i_ that could control pigmentation gene expression.

To confirm the involvement of p300/CBP in pigmentation, we incubated the B16 cells with C646 a selective inhibitor of the p300/CBP HAT activity. Upon inhibition, B16 cells despite being subjected to pigmentation permissive 5% CO_2_ condition did not pigment **(Fig 5H)**. The expression of melanocyte differentiation genes was downregulated with marginal changes in MITF transcript levels, confirming that p300/CBP facilitate the melanocyte maturation process by enhancing the expression of differentiation genes. Taken together, we demonstrate that MITF activates Ca14 transcriptionally; Ca14 increases pH_i_ that in turn can activate p300 HAT activity, which acetylates histone H3 in the promoters of melanocyte differentiation genes facilitating their transcriptional activation by MITF (**Fig 5I**). Hence this feed forward loop enables quick and heightened expression of differentiation genes under conditions where the cells require rapid pigmentation.

### Targeted mutation of ca14 reiterates the role of feed forward loop on pigmentation

We chose to address the role of this feed forward loop mediated by CA14 in pigmentation under physiological conditions wherein rapid and programmed induction of pigmentation genes is involved. We created a germline mutant in zebrafish by targeting the ca14 coding region using the CRISPR-Cas9 system. We injected Cas9 - guide RNA complex to fertilized zebrafish embryos at the single cell stage. We observed a variety of phenotypes in F0, which include microcephaly, small eye size, mild enlargement of heart and decreased pigmentation. These phenotypes recapitulate the known high expression of Ca14 in brain, heart and eye. After screening for mutants using T7 endonuclease assay, siblings were grown to adulthood. Genotyping revealed a frame-shift mutation at the third codon by the deletion of two bases. Thereby the mutant gene would encode a truncated protein lacking most the coding amino acids (**Supplementary Fig S8**). Further experiments were carried out using F2 embryos obtained from the in-cross of a homozygous mutant male and a heterozygous female fish with the same mutation. We obtained around 50% of embryos with a smaller eye size and decreased pigmentation, following a Mendelian pattern of inheritance. Mutant embryos showed the two base deletion and the non-phenotypic siblings were heterozygous based on PCR confirmation of the mutation. Thereby the pigmentation phenotype observed in the morphant embryos is recapitulated in the genetic mutants, hence ascertaining the regulatory role of Ca14 in the maturation of melanocytes (**Fig 6A**).

**Fig 6:**
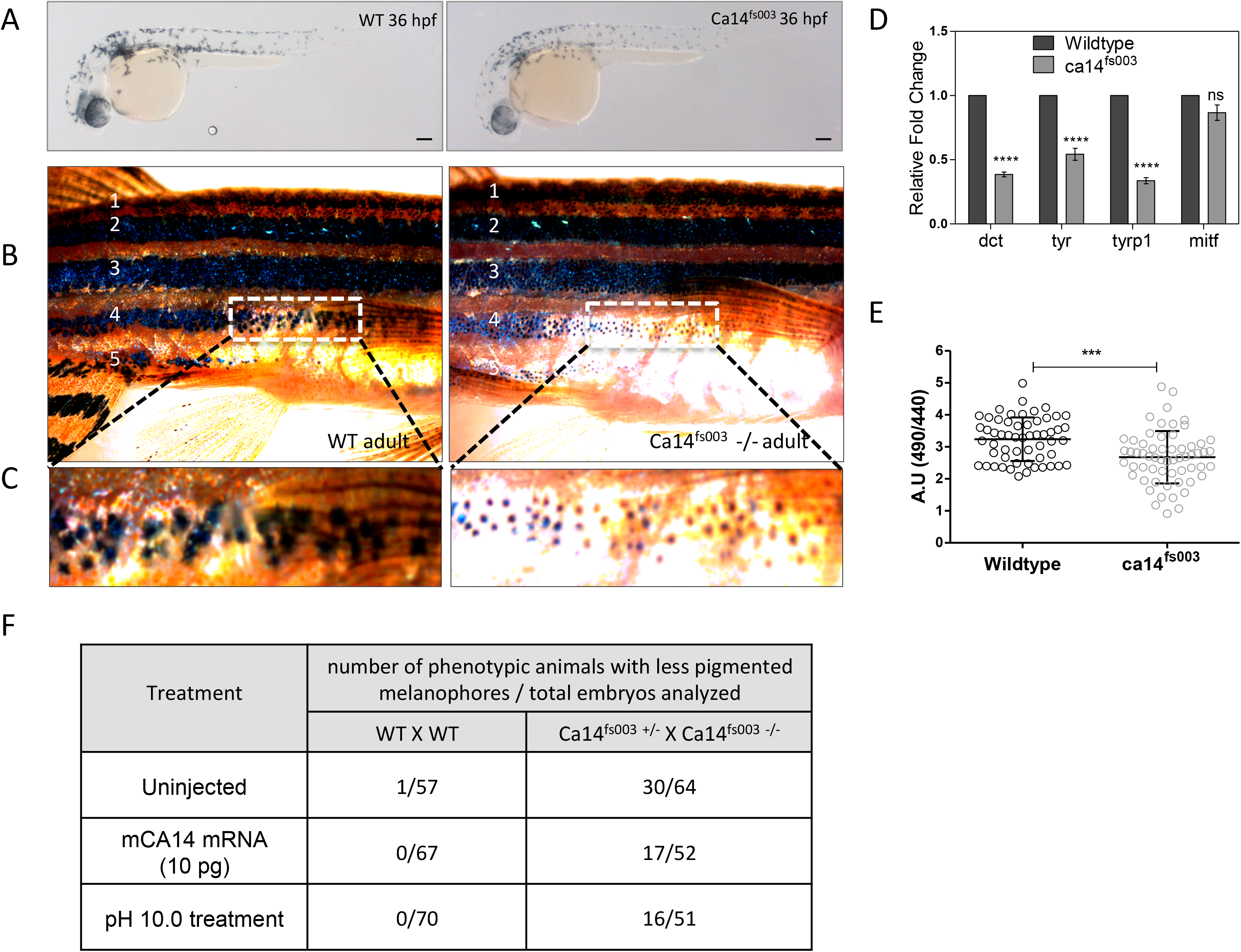
Tragetted null mutation of Ca14 by CRISPR demonstrates immature acidic melanocytes. A.Brightfield images of the lateral view of CRISPR targeted mutant *ca14*^*fs003*^ and control embryos at 36 hours post fertilization (hpf) in F2 generation fishes. Scale bar 100 μm. B.Wildtype and *ca14* ^*fs003*-/-^ adult male animals were dark adapted for 24h. Image of the lateral view was captured under identical lighting and image capture settings. C.Zoomed up portion from the fourth lateral line demonstrates melanophores to be laden with less melanin content in the mutant animal. D.qRT-PCR quantification of pigmentation genes *tyrosinase* (*tyr*), *dopachrome tautomerase* (*dct*), *tyrosinase related protein 1b* (*tyrp1b*) *and micropthalmia associated factor a (mitfa)* between control and *ca14* ^*fs003*^ at 36 hpf, using *rps11* gene as the normalization reference. While mitfa remains unaffected all other pigmentation genes are downregulated in the absence of functional ca14. E.Intracellular pH_i_ probed by BCECF-AM in *ca14* **^*fs003*^** and sibling control zebrafish embryos. F.Readings were obtained from trunk region melanophores. n = 3 expts, at least 10 embryos each. A decrease in the ratio indicates acidification of melanocytes. G.Wild type as well as embryos obtained from the cross of *ca14* **^*fs003*-/-^** and *ca14* **^*fs003*+/-^** were left uninjected or treated with embyo water buffered at pH 10.0 (between 18hpf till 36 hpf) or injected with 10 pg of in vitro transcribed mouse Ca14 mRNA. Embryos were scored for pigmentation at 36 hpf. Ratios represent number of less pigmented phenotypic embroys to the total number of embryos scored. The proporition of less pigmented animals decreased upon mouse Ca14 mRNA injection as well as increased extracellular alkalinization.

In the adult stage Ca14 mutation had a visible decrease in pigmentation, however high melanophore density present in the lateral and dorsal region precluded assessment of melanin. We therefore subjected the wild type adult and the *ca14*^*fs003*-/-^ fishes to dark adaptation to disperse the existing melanin within the melanocyte so that the content could be easily assessed. We observed distinct non-overlapping melanophores present in fourth and fifth stripes near the caudal fin and in this region a substantial reduction in the melanin content could be observed in the mutant fish (**Fig 6B & C**).

We chose 36 hpf time point to analyze the expression of differentiation genes when the pigment cells undergo migration and maturation. A decreased gene expression of *tyr, tyrp1b* and *dct* could be observed in the mutant embryos (**Fig 6D**); confirming that the cells are in an immature less pigmented state and the pigmentation promoting gene expression is severely reduced in the absence of Ca14.

### Ca14 mutant melanophores are acidic with reduced pigmentation

We then subjected the wild type and the *ca14*^*fs003*-/-^ to intracellular pH measurements. Mutant embryos had lower ratiometric values of BCECF (490/440) suggesting an acidic intracellular pH in melanocytes (**Fig 6E**). We also attempted a chemical rescue approach by subjecting the mutant embryos to embryo water buffered at various pH between 24 hpf to 36 hpf, when the melanocyte maturation commences. While the acidic pH 5 did not show a rescue rather showed a marginal increase in deformities, alkaline pH 10 rescued the phenotype marginally and animals with small eye had now substantially high melanin content (**Fig 6F**). The extent of pH-mediated rescue was comparable when the animals were injected with mouse Ca14 mRNA. It is interesting to note that; in the *ca14*^*fs003*-/-^ mutant number of melanocytes remains unaltered but the expression of differentiation effectors is relatively low. Therefore in the absence of ca14, melanocytes would still respond to external cues and activate Mitf, however the extent of pigmentation would be severely curtailed. The mutant melanophores clearly showed decreased melanin content, thereby unequivocally establishing the role of the feed forward loop involving MITF, Ca14 and the differentiation genes in ensuring a heightened pigmentation.

## Discussion

Several genes that alter melanin content and melanocyte functions are easily identified using naturally occurring mutants and targeted gene disruptions (Bennett & Lamoreux, 2003). Recently, siRNA based genome-wide screen revealed some of the important pigmentation genes (Ganesan et al, 2008). While these perturbation-based approaches reveal the identity of components, biologically relevant regulators and the physiological context are often the most difficult to decipher. In our earlier work, *ca14* gene was observed to co-regulate with the pigmentation status in B16 melanoma cells (Natarajan et al, 2014). Taking clue from this observation and the indication that *ca14* could be under the control of Mitf, we have embarked on identifying the role of CA14 in melanocyte functions. In the current study, using cell as well as whole animal based approach using zebrafish, we demonstrate an important role for CA14 and the downstream pH changes as an amplifier of the maturation process in melanocytes.

As a general theme pH changes and cell fate decisions have attracted a lot of attention, but precise mechanistic understanding is often limited by the widespread changes brought about by pH (Tatapudy et al, 2017). CA14 was identified as a potential Mitf target gene based on promoter binding and up-regulation of mRNA in melanocytes overexpressing Mitf (Hoek et al, 2008; Strub et al, 2011). However the physiological implications of this observation was not readily apparent. Based on our study, we propose a model wherein CA14 is an early gene induced by Mitf, and together the two accelerate the gene expression of a subset of pigmentation genes resulting in a feed-forward amplification. This mode of network interaction is seen often during cell differentiation programs. For instance, during chondrogenic differentiation two key transcription factors Bmp2 and Shh, operate to regulate Sox9 in a positive feedback loop to stimulate cellular differentiation (Zeng et al, 2002).

While the link between pH and cell differentiation is often observed across many systems, mechanisms that could connect changes in pH to pigmentation gene expression were not immediately apparent. An increase in intracellular pH (pH_i_) is necessary for the differentiation of follicle stem cells in drosophila (FSCs) as well as mouse embryonic stem cells (mESCs) (Ulmschneider et al, 2016). *Drosophila* Na^+^–H^+^ exchanger *DNhe2* is involved in lowering the pH_i_ in differentiating cells and impairs prefollicular cell differentiation, whereas increasing pH_i_ promotes excess differentiation toward a polar/stalk cell fate through the suppression of Hedgehog pathway. Together with our observations, it is now apparent that this metabolic link involving pH and cell differentiation may be a far more widespread mechanism that would be operational in many cell types. Earlier studies indicate that Mitf regulates differentiation of precursors to mature osteoclasts by the induction of carbonic anhydrase II, which is responsible for extracellular acidification (Lehenkari et al, 1998; Lu et al, 2010). It is likely that pH-mediated induction of differentiation and its regulation by Mitf may be an underlying theme conserved across several cell types.

The role of p300/CBP in MITF mediated gene expression has emerged from multiple evidences. Activated MITF physically interacts with p300/CBP, and Mitf immunoprecipitate has associated HAT acitivity (Price et al, 1998) (Sato et al, 1997). However the physiological effect of this association remained enigmatic, as MITF mutants defective in binding to p300/CBP still activated transcription (Vachtenheim et al, 2007). The footprint of a SWI/SNF complex involving BRG1 on MITF targets has distinct clusters of genes with high as well as low H3K27 acetylation status (Laurette et al, 2015). Thereby it is likely that distinct subsets of MITF targets have different levels of activation. This is also recently alluded to by (Malcov-Brog et al, 2018). The authors demonstrate that H3K27 acetylation pattern is unusually high in pigmentation related MITF targets compared to proliferation and sustenance genes. Hence making dynamic regulation of H3K27 acetylation a prerequisite for a subset of genes, while others are less dependent on this activation. The identification of Ca14 mediated p300 activation elucidated in this study provides a plausible mechanism for this crucial but so far intriguing observation on the context dependent target selectivity by MITF.

p300 histone acetyl transferase is a central epigenetic regulator and the observed pH mediated activation is likely to be a general phenomenon. The crystal structure of the p300 HAT domain revealed an unusual hit and run catalytic “Theorell-Chance” mechanism (Liu et al, 2008). Subsequent molecular dynamic simulations suggest a proton relay from the ε amino group of the acceptor lysine substrate and it is predicted that the involved side chains have a high pK_a_ (Zhang et al, 2014). The standard enzymatic assays involving p300 are routinely carried out at an alkaline pH 7.8 - 8.0 which would favour deprotonation providing a molecular mechanism of pH-mediated HAT activation. Thereby the transient increase in pH mediated by Ca14 facilitates H3K27 acetylation by p300 thereby rendering activation of these genes.

Ca14 mediated feed-forward loop would be operational under conditions wherein rapid melanocyte maturation is required. This is anticipated during UV induced sun tanning response as well as developmental states wherein melanocytes mature into a high melanin containing cells. Recent study identified *ca14* to be downregulated in the lesional depigmented skin of vitiligo subjects (Yu et al, 2012). It is tempting to speculate that the depigmentation in vitiligo could be contributed by the decrease in CA14 that could affect melanocyte maturation, in addition to the loss of mature melanocytes. Advances made in this study are therefore of immense relevance in the recently emerging role of biochemical milieu as an intermediate effector in determining the cell fate decisions.

## Materials and Methods

### Cell line and culture

B16 mouse melanoma cell line was cultured in DMEM-high glucose (Gibco, Life Technologies) medium supplemented with 10% fetal bovine serum (FBS; Gibco, Life Technologies) and cells were maintained in 5% (or at 10% CO_2_ when indicated) in a 37°C incubator. The orthologous non-cancerous murine melanocyte line Melan-A was cultured at10% CO_2_ at 37° C in RPMI-1640 medium (Gibco, Life Technologies) supplemented with 10% fetal bovine serum (FBS; Gibco, Life Technologies), 400nM Phorbol-Myrystyl-13-acetate (PMA; Sigma) and 0.003% Phenylthiourea (PTU, Sigma). Primary human melanocytes were grown in proliferative conditions in PMA containing medium MBM4 (Lonza). For differentiation cells were switched to M254 medium for 3-4 population doublings (ThermoFisher Scientific, Life Technologies).

### Setup of pigmentation oscillation model in B16 cells

Detailed Protocol described in (Natarajan et al, 2014). Briefly, B16 cells were seeded at density of 100 cells/cm^2^ in DMEM-high glucose medium supplemented with 10% FBS. Cells were cultured at 5% CO_2_ and were harvested at day 4, 6, and 8 for downstream experiments. The cells progressively pigmented and the expression of pigmentation genes were induced in this model system (**Supplementary Fig S2**). To modulate the pH cells were alternatively cultured in 10% CO_2_ and subsequent changes in pH, melanin content and gene expression were investigated. P300/CBP inhibitor C646 or vehicle control DMSO treatment was performed at Day 1 of pigmentation oscillator and terminated on Day 6 for downstream analysis.

### *Generation of ca14 promoter construct*, Ca14 *expression construct and the inactive mutant*

The *ca14* 3kb promoter (3000 bp upstream of transcription start site) was amplified from mouse genomic DNA and cloned upstream of luciferase cassette in KpnI/HindIII site of pGL4.23 (Promega) using primers listed in **Supplementary table 1**. Mouse Ca14 coding sequence was amplified from mouse B16 cDNA and cloned in mcherryN1 vector (Clontech) in KpnI/HindIII site. Site directed mutagenesis for CA14 (Thr199Ile) was carried out using SDM II kit (Agilent) using primers listed in **Supplementary table 1**. Truncated constructs of 1kb (proximal, middle, distal from TSS) were generated from *Ca14* 3kb promoter. *Dct* promoter used in this study is reported elsewhere (Natarajan et al, 2014).

### Measurement of intracellular pH

#### For mammalian cells

Intracellular pH (pH_i_) was measured by using the ratiometric dye BCECF-AM (Molecular probes, Thermo scientific), as described in (Natarajan et al, 2014). Briefly cells were plated in 35 mm dishes (IBIDI) and on the day of pH measurement cells were incubated with 0.2µM of BCECF-AM dye for 2min at 37#x00B0;C and 5% CO_2_. In order to generate a pH calibration curve, cells were incubated in pH calibration solution (in mM:1Glucose, 140KCl, 1MgSO_4_, 30HEPES, 25NaCl, 1CaCl_2_, 1NaH_2_PO_4_)with pH range of 6– 8.5 and added 10µMNigericin (Thermofisher Scientific). Quantitative images were acquired and fluorescence ratio was measured at dual excitation wavelength of 440 and 490 nm in Leica SP8 confocal microscope.

#### For zebrafish embryos

2 dpf zebrafish embryos were taken and dechorinated. 10 embryos were incubated with 10mM BCECF-AM (Thermofisher scientific) in embryo water at 28C incubator for 15 minutes. After incubation the embryos were washed, embedded in methylcellulose and taken for quantitative imaging using Leica SP8 confocal microscope.

### Plasmid and silencing RNA Transfections

B16 cells were trypsinized and seeded at density of 1X10^5^ cells/well in 6 well plate (BD Bioscience) and incubated overnight in antibiotic containing DMEM + 10% FBS. At the time of transfection, the cells were replaced with serum and antibiotic free media OptiMEM (Gibco, Life Technologies). Lipofectamine 2000 was used at a ratio of 1:2 with CA14 siRNA or the scrambled control siRNA (Dharmacon). The cells were incubated for 6 hours with the transfection mixture containing Lipofectamine 2000, OptiMEM and the siRNA. The media was then replaced with antibiotic containing DMEM+10% FBS and incubated for 72 hours and downstream experiments were performed.

NHEM cells were trypsinized and seeded at density of 1X10^5^cells/well in 6-well plate (BD Bioscience) and incubated overnight with antibiotic containing M254 (Thermofisher Scientific). Next day cells were washed with DPBS (Gibco, Life Technologies) and were replaced with antibiotic free OptiMEM media (Gibco, Life Technologies). Cellfectin (Invitrogen) was used at a ratio of 1:2 with MITF siRNA and CA14 siRNA (Dharmacon) and incubated for 6 hours. The transfection mixture was replaced with antibiotic containing M254 media and incubated for 72 hours.

### Treatment for the induction of MITF

Melan-a cells were trypsinized and seeded at density of 1X10^5^ cells/well in 6 well plate (BD Bioscience) and incubated overnight with antibiotic containing RPMI 1640+10% FBS. Next day fresh media was added to the cells and treatment was performed with α-MSH (1000nM; Sigma) and IBMX (60uM; Sigma). The cells were incubated at 37° C at 10% CO_2_ and were then harvested for protein isolation after 48 and 72 hours.

### RNA Isolation & cDNA synthesis

Cells were trypsinised with 0.1% trypsin and the pellet was washed twice with 1X Phosphate Buffered Saline (PBS; Gibco Life Technologies). To the pelleted cells 1ml of TriZol (Ambion, Invitrogen) was added and stored at −80° C overnight. The RNA was isolated using standard Trizol based method. The isolated RNA was subjected to DNAse (Qiagen) treatment for 20 minutes at room temperature. Column purification of the RNA was performed using RNAeasy mini kit (Qiagen, Cat 74104) according to manufacturer’s protocol. 100-500 ng of RNA was taken for cDNA synthesis using Superscript III (Invitrogen, Life Technologies) according to manufacturer’s protocol. Q-RT pcr experiments were performed either by SYBR green or TaqMAN assay probes (Details provided in **Supplementary table 1**) according to manufacturer’s protocol using Lightcycler 480 II (Roche).

### Isolation of nuclear and melanosomal fractions

Nuclear proteins were extracted using NE-PER^™^ nuclear and cytoplasmic extraction reagents (Thermoscientific; 78833) according to manufacturer’s protocol. Melanosomes were isolated using protocol previously described (Watabe et al, 2005). The B16 tumor from mice was excised and washed in homogenization buffer and was homogenized with 120 strokes of dounce glass homogenizer. Post nuclear supernatant was isolated and then separated on a stepwise density gradient and centrifuged at 1,00,000 g for 1 hour at 4 ° C in a swing out rotor. The stage III and IV melanosomes which preferentially localized to the highest density fraction (1.8-2.0) were collected and protein was isolated using NP40 lysis buffer.

### Protein Isolation and Western Blot analysis

Cells were trypsinised with 0.1% trypsin and the pellet was washed twice with 1X Phosphate Buffered Saline (PBS; Gibco Life Technologies). NP40 lysis buffer (Invitrogen) was added to the pellet and was incubated on ice for 30 minutes with pipetting at interval of 10 minutes. The cells were centrifuged down at 13,000 rpm for 30 minutes at 4° C (Eppendorf Centrifuge 5415 R). The supernatant was collected and transferred to a fresh microfuge tube. For Histone blots lamelli lysis buffer was used. The protein was estimated using standard BCA protocol (Pierce BCA protein assay kit; Thermoscientific). Equal amount of protein from each sample were resolved in 10 or 12% % SDS gel in 1X Tris glycine buffer. The gel was blotted onto 0.45 uM PVDF membrane (Millipore) at 300mA for 2 hours. 5% Skim milk was used for blocking for 1 hour at room temperature. Incubation with primary abtibody was performed for overnight at 4° C. Primary antibody dilutions are provided in **Supplementary table 1**. After washing the blot with 1X TBST, the blot was incubated with HRP conjugated secondary antibodyfor 1 hour at room temperature. After washing with 1X TBST the blot was developed using Immobilon Western (Millipore) in the Syngene GBOX Chemiluminescence instrument. Densitometry analysis was performed using ImageJ software.

### Chromatin Immunoprecipitation and q-RT PCR

B16 Melanoma cells at 80% confluence was fixed with 10% formalin (Sigma Cat No HT501128) and incubated at 37 C for 10 minutes. 2.5 M Glycine was added to the cells and again incubated at 37 C for 10 minutes. Cells were washed with Ice cold 1X PBS containing protease inhibitors. Cells were then scraped and centrifuged at 1000 rpm for 5 minutes at 4C. The cell pellet was lysed in SDS lysis buffer (1% SDS, 10mM EDTA, 50mM TRIS (pH 8.1)) on ice for 30 minutes. The cells were then sonicated on bioruptor (DIAGENODE) in ice. The chromatin lysate was then estimated for protein content using BCA kit (Pierce). 5ug of MITF C5 antibody (Abcam) was taken and incubated with Protein G Dynabeads (Thermoscientific) overnight at 4 C on a rotator. 3ug of acetylated H3K27 antibody was incubated with Protein A dynabeads (Thermoscientific) overnight at 4 C on a rotator. The next day the sera was cleared and washed with Ice cold dilution buffer (2mM EDTA, 150mM Nacl, 20mM Tris HCl (pH 8)). 500 ug of chromatin lysate was added to the beads and the final volume was made up to 750ul using the dilution buffer and incubated for 6 hrs at 4C in a rotator. 10% of the lysate was kept separately as input. After incubation the magnetic beads were washed with Low salt Buffer (0.1 % SDS, 1% Triton X 100, 2mM EDTA, 20mM Tris HCl (pH 8), 150mM NaCl), high salt buffer (0.1 % SDS, 1% Triton X 100, 2mM EDTA, 20mM Tris HCl (pH 8), 500mM NaCl) and LiCl buffer (0.25 M LiCl, 1% Igepal C-630, 1mM EDTA, 10mM Tris HCl (pH 8), 1% deoxycholate). Finally the magnetic beads were incubated overnight at 65 C in elution buffer (1% SDS, 0.75% sodium bicarbonate) and 1ul of 20mg/ml proteinase K (Sigma) for elution and subsequent reverse crosslinking. The magnetic beads were separated from supernatant and column purified using QIAgen pcr purification kit, the input control was also included in the purification step. SYBR qRT PCR was setup using 5ul of eluted DNA and graphs were plotted as percentage input.

### Immunofluorescence

The cells plated on coverslips were washed twice with 1X PBS and then fixed with 4% PFA at 37° C for 20 minutes. The cells were again washed twice with 1X PBS and permeabilized with 0.01 % Triton X100 (Sigma). The cells were blocked with 5% Normal goat serum (Jackson Laboratories) overnight at 4° C. The cells were washed twice with PBST (1X PBS with 0.01% Tween-20). The cells were then incubated with 1:50 dilution of CA14 antibody (Abcam) in a moist chamber for 1 h at room temperature. Incubation with secondary antibody (Alexafluor488; Molecular probes, Thermoscientific) was performed at room temperature for 1 h. The cells were again washed with PBST. The cells were then mounted on slides using Antifade slowfade DAPI (Molecular probes, Invitrogen) and visualized using Confocal Microscope (Zeiss LSM 510 or Leica SP8).

### P300 HAT activity assay

HAT activity assay was performed using KAT3B/P300 inhibitor fluorometric assay screening kit (Abcam, ab196996) according to manufacturers protocol with minor modification. Tris-Cl pH 7.15 and pH 7.95 was added at an additional concentration of 50mM to buffer provided in the kit and the assays were performed with the modified buffers.

### Zebrafish Ethics Statement

Fish experiments were performed in strict accordance with the recommendations and guidelines laid down by the CSIR Institute of Genomics and Integrative Biology, India. The protocol was approved by the Institutional Animal Ethics Committee (IAEC) of the CSIR Institute of Genomics and Integrative Biology, India (Proposal No 45a). All efforts were made to minimize animal suffering.

### Zebrafish strains and maintenance

The wild type strain Assam WT (ASWT) and Tyrp1 reporter, *Tg(ftyrp1: GFP)* zebrafish lines were used for morpholino injections and were maintained according to standard zebrafish husbandry protocols. ftyrp1:GFP plasmid was a kind gift from Dr Xiangyun Wei (Zou et al, 2006), University of Pittsburgh School of Medicine and the transgenic line was created at CSIR-IGIB zebrafish facility using tol2 transposase microinjections in ASWT line. Phenylthiourea (PTU; 0.003%) was added to embryo water before 24 hours post fertilization (hpf) to prevent melanin from masking the GFP fluorescence. For pH rescue experiments the embryos were grown in embryo media buffered with 10mM HEPES.

### Morpholino knockdown of zebrafish Ca14

Antisense morpholino was synthesized from Gene Tools against Ca14 of zebrafish. The morpholino was designed to block the translation of car14 gene. Morpholino sequence with translation initiation site (initiator ATG codon) is underlined CC<u>ATG</u>ATTTCACTATTCTCCCTACA. Standard control was obtained from Genetools and injected at same dosage as ca14 morphlino.

### ca14 CRISPR mutant generation

Zebrafish ca14 CRISPR was designed using ECRISP software (http://www.e-crisp.org/E-CRISP/). The CRISPR sgRNA sequence was in vitro transcribed using mMessage mMachine T7 ULTRA kit (Thermofisher scientific). 300 pg of sgRNA was injected along with 500 pg of spcas9 protein (kind gift from Dr Souvik Maiti, CSIR-IGIB). The F0 embryos were screened using T7 endonuclease assay for mutations and the putative mutant siblings were grown to adulthood and inbred to get F1 animals. Mutations in the F1 fishes were confirmed using sanger sequencing.

The cross involving the ca14fs003 reported herein involves a *ca14^fs003^-/-* male with a *ca14^fs003^+/-* female that accounts for 50% of the embryos to be phenotypic. The genotypes of the phenotypic embryos and normally pigmented embryos arising from this cross were ascertained to be *ca14^fs003^-/-* and *ca14^fs003^+/-* respectively using PCR based amplification.

### Zebrafish imaging

The embryos were manually dechorinated and embedded in 2% methylcellulose. Control and Morphant embryos were imaged laterally and dorsally for melanophore quantitation. Brightfield Images were taken using Zeiss Stemi 2000-C microscope; for fluorescence microscopy Zeiss Axioscope A1 was used.

### RNA Isolation from embryos

Total RNA was extracted from zebrafish embryos using TriZol (Invitrogen). cDNA was synthesized using Superscript III kit (Invitrogen). Real time quantitative PCR was performed using SYBR Green (Kapa Biosystems) or TAQMAN probes (details provided in **supplementary table 1**) and data was generated in ROCHE Lightcycler 480 II.

### mRNA injection in zebrafish embryos

For rescue experiments the coding sequence of mouse *Ca14* gene was cloned with kozak sequence inserted in front of translation start site **(Supplementary table 1)**. The amplicon was cloned into TOPO-Zero blunt vector (Thermofischer Scientific). RNA was made using *in-vitro* transcription kit T7 Ultra mRNA synthesis kit (Ambio – Thermofischer Scientific). 10 pg of Ca14 WT or Ca14_T199I_ RNA was injected into the cell at one cell stage Ca14^fs003^ embryos.

### Estimation of Melanin content

Dorsal images of Control and ca14 morphant embryos were taken at 2 dpf. The images were imported to ImageJ and mean grey values were taken for melanophores of each embryo set. Mean grey values are inverse proportional to the melanin content of the cell. Coresponding values were then plotted using Graphpad Prism.

### Statistical analysis and Graphs

Student’s t test was performed to obtain statistical significance in the data. Asterisk on the error bar corresponds to *(P ≤ 0.05), ** (P ≤ 0.01), *** (P ≤ 0.001), **** (P ≤ 0.0001) and ns (P > 0.05). Graphs were plotted using Graphpad prism.

## Acknowledgement

This work was supported by the Council for Scientific and Industrial Research (CSIR), India through grant (TOUCH-BSC0302) and Department of Biotechnology through the grant (GAP0182). We acknowledge the infrastructural support of CSIR to the imaging facility (VISION-BSC0403).

## Author contributions

D.A.R, V.G, R.S.G and T.N.V designed experiments. D.A.R, V.G, F.S and Y.J.S, executed the experiments with cultured cells. A.S performed experiments pertaining to electron microscopy. D.A.R, S.S and T.N.V were involved in the design and execution of zebrafish experiments. D.A.R, Y.J.S, V.G along with S.S, R.S.G and T.N.V were involved in data analysis, interpretations and writing of the manuscript.

## Supplementary Information

### SUPPLEMENTARY FIGURE LEGENDS

**Fig S1:**
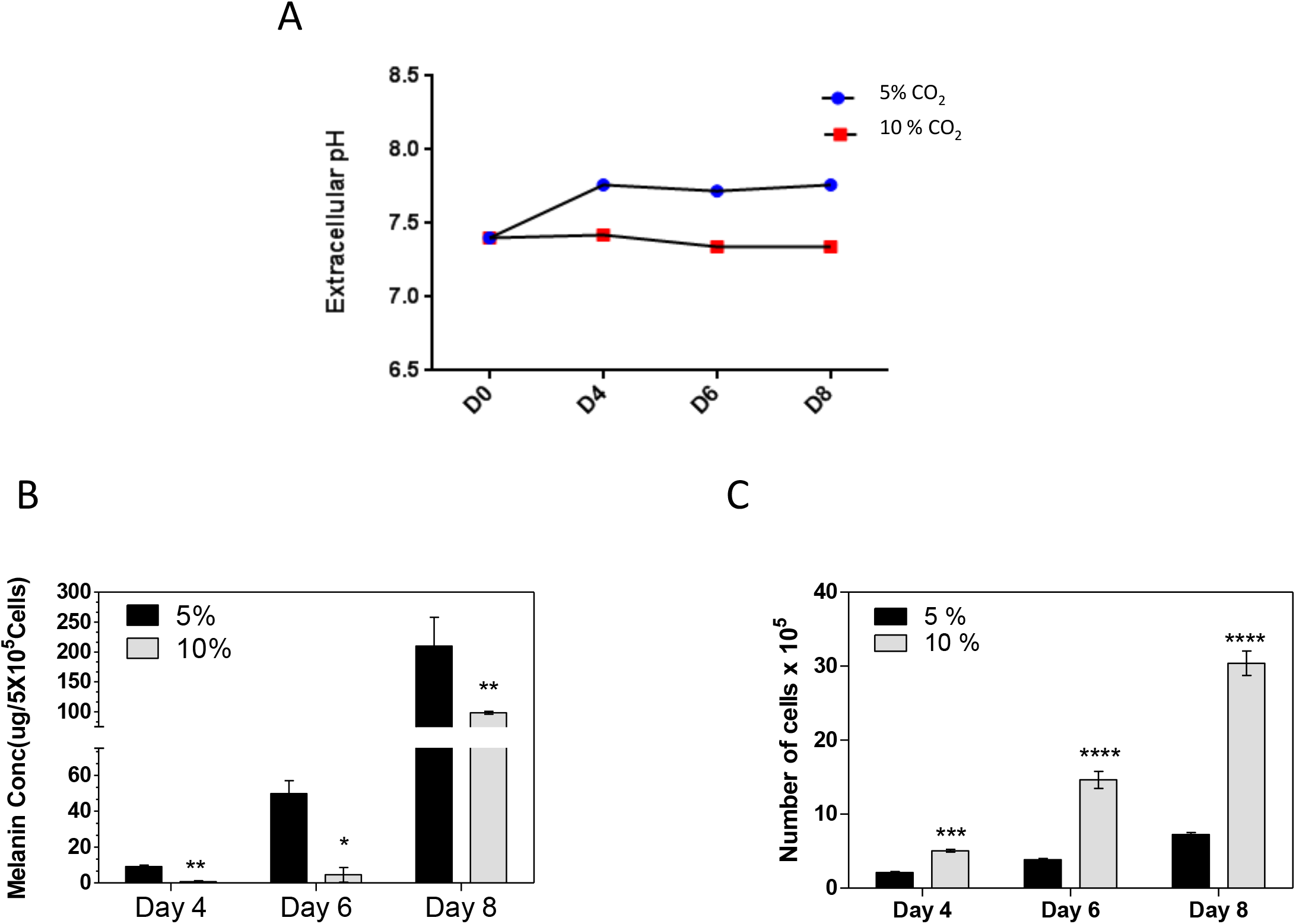
10% CO_2_ leads to extracellular acidification leading to decreased melanin content and increased proliferation. A.Line graph showing extracellular pH at various days of pigmentation in 5% and 10% CO_2_ cultured B16 cells. B.Bar graph depicts the colorimetric melanin estimation of B16 melanoma cells cultured under 5% CO_2_ and 10% CO_2_ growth conditions for indicated days using synthetic melanin as a standard. Error bars represents independent biological replicate, n=3. C.Alteration of cellular pH by the modulation of CO_2_ results in changes in cell proliferation in B16 cells as measured by cell count.

**Fig S2:**
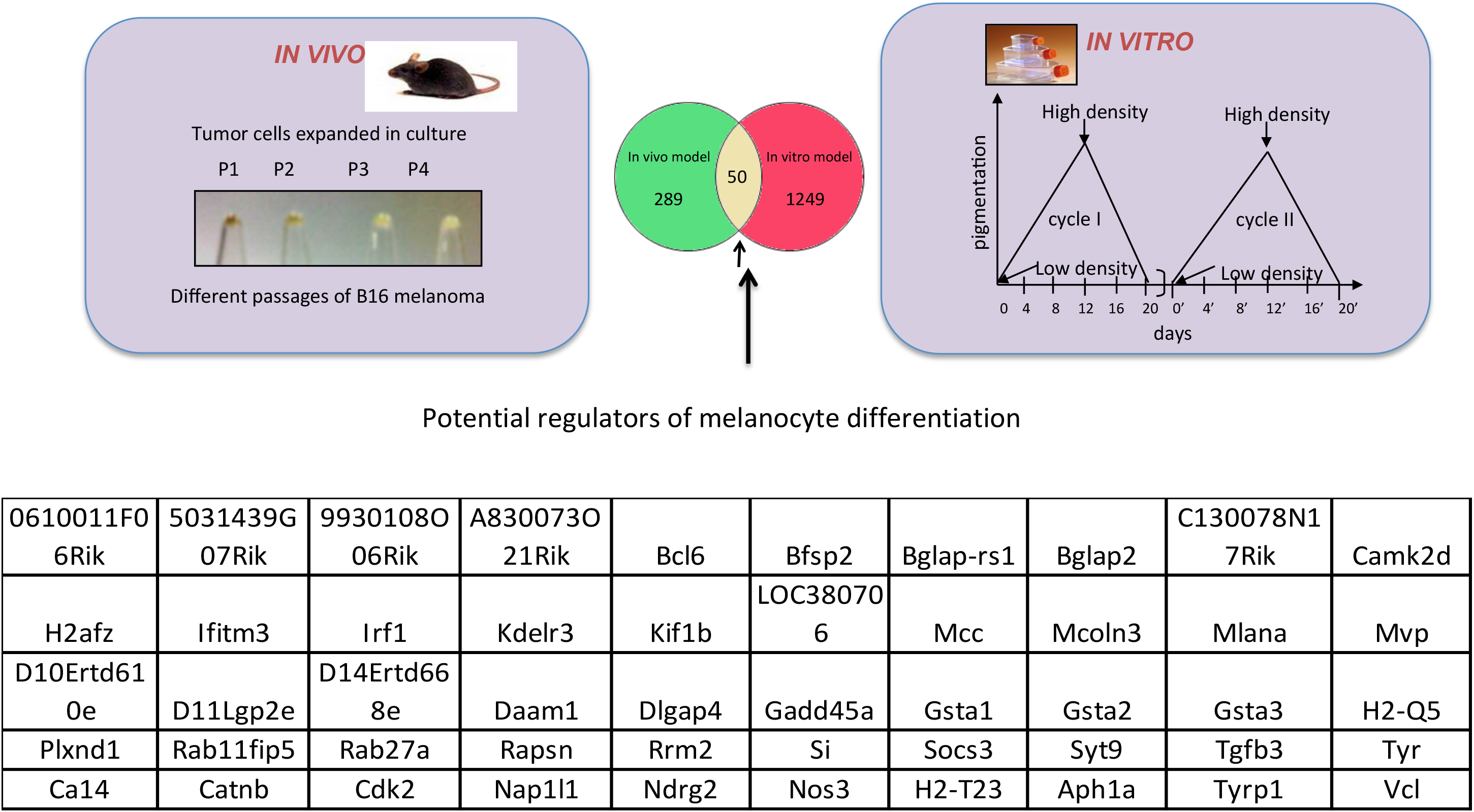
Gene expression data analysis from pigmentation models to identify putative effectors of melanocyte differentiation. B16 melanoma cells were made to transit between pigmented (differentiated) state and depigmented state state, using two methods: by passaging the depigmented cells in mouse as a tumor and culturing the cells in vitro. Microarray was performed on the four different *in vitro* passages (P1 to P4). Depigmented B16 cells were cultured *in vitro* at a low density and allowed to pigment for 12 days and to depigment for the next 8 days for two cycles. Microarray was carried out from Day 0 depigmented Day 4, Day 8 and Day 12 (pigmented) and Day 16 and Day 20 (depigmented). Expression profiles of genes that showed a concomitant pattern of regulation across both models of melanocyte differentiation is listed. (GSE54359).

**Fig S3:**
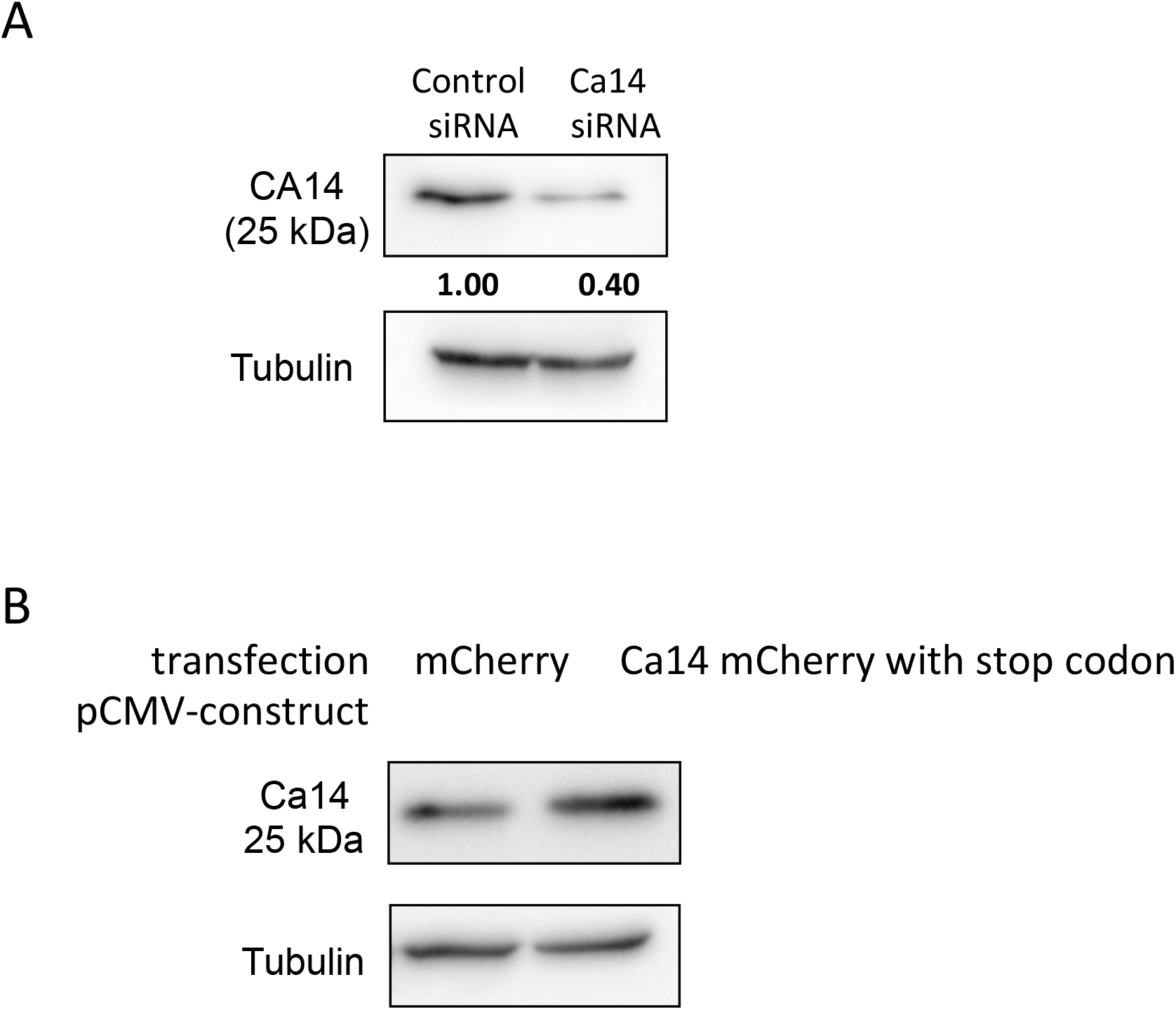
Validation of Ca14 antibody used in the study. A.Blots depicting the levels of Ca14 protein upon Ca14 siRNA mediated knockdown. Tubulin was used as normalizing control. B.Blots depicting the levels of Ca14 protein upon overexpression. Tubulin was used as normalizing control.

**Fig S4:**
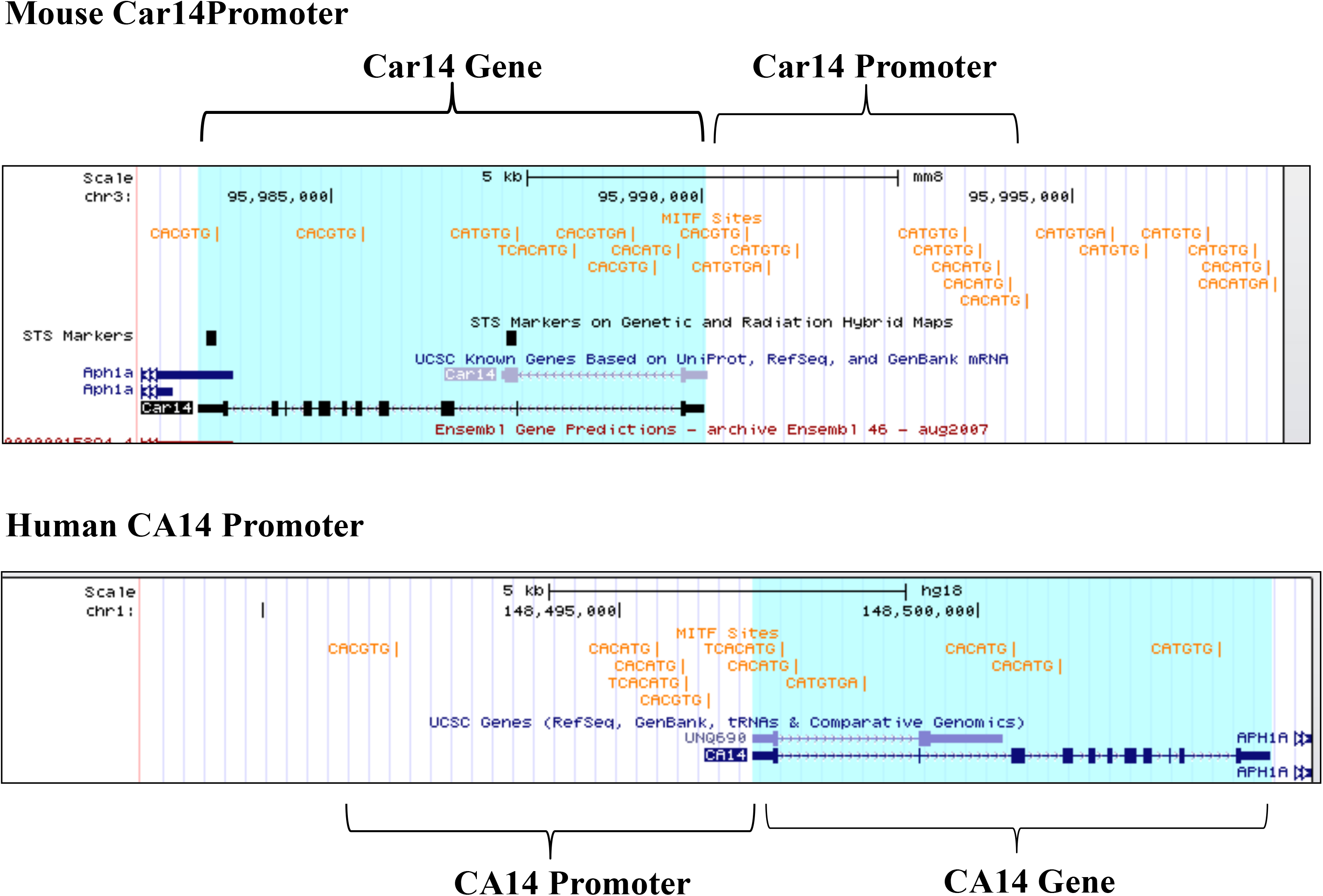
M box sequences in Human and Mouse ca14 promoter and genic regions.

**Fig S5:**
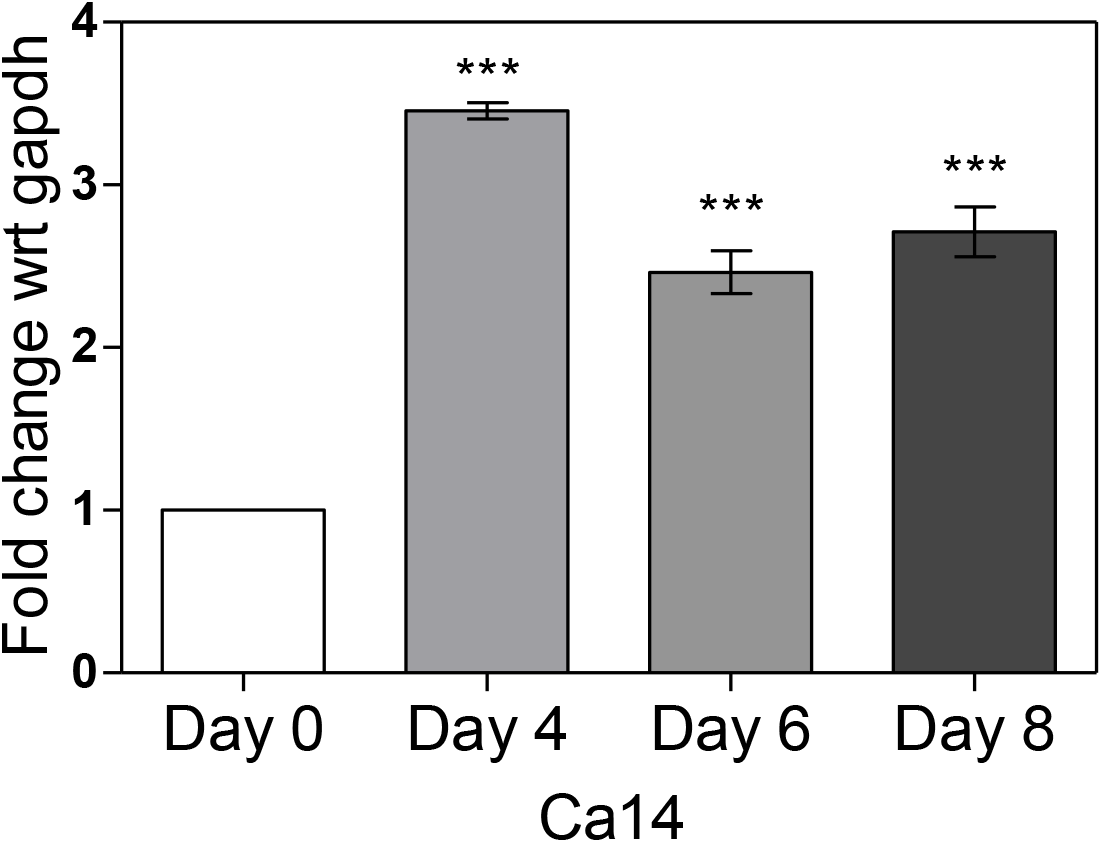
mRNA expression of carbonic anhydrase 14 during pigmentation. Bar graph depicting the relative mRNA levels of Ca14 during pigmentation in mouse B16 melanoma cells. Error bars represent SEM from biological replicates across three different experiments.

**Fig S6:**
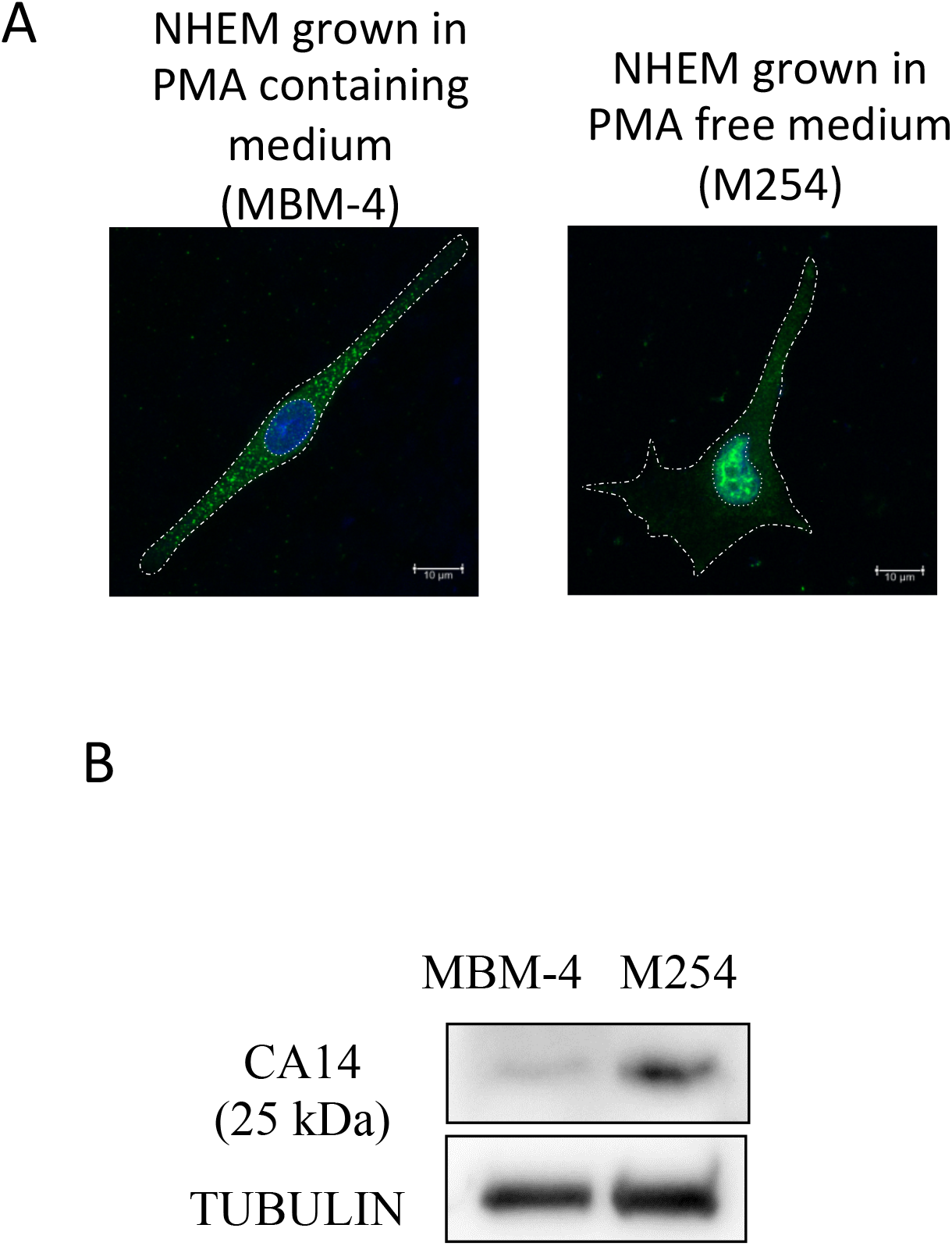
Changes in CA14 expression correlate with pigmentation status of primary human melanocytes. A.Confocal image of Normal Human Epidermal Melanocytes (NHEM) cultured under different media conditions using MBM4 and M254 media available commercially. B16 cells were stained using for CA14 antibody (green) and the nucleus is counter stained using DAPI (blue). B.Protein levels of CA14 probed by western blot analysis in NHEM grown under different media conditions that retain melanocytes in a proliferative condition (MBM-4 medium with phorbol myristyl acetate (PMA)) and M254 (without PMA) that facilitates pigmentation.

**Fig S7:**
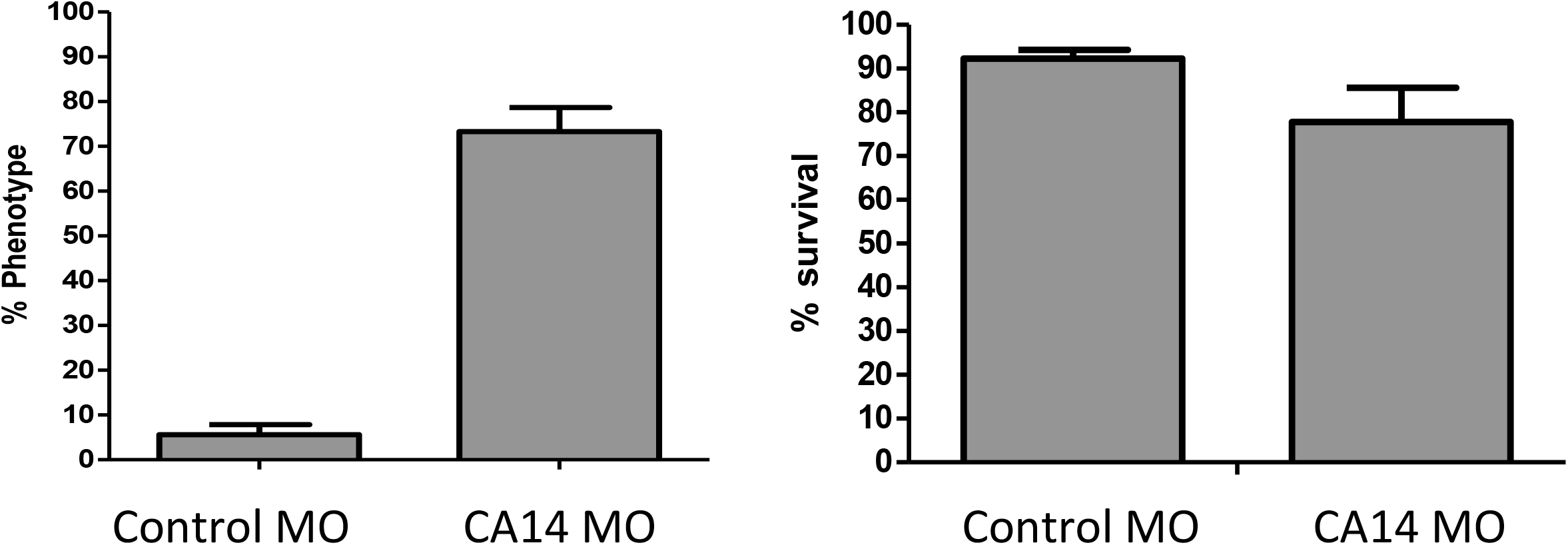
ca14 morpholino causes pigmentation phenotype without affecting the survival of embryos. A.Percentage phenotype in Ca14 morpholino injected embryos w.r.t control morpholino represented as bar graph. Error bars represent SEM across four independent experiments. B.Percentage survival in Ca14 morpholino injected embryos w.r.t control morpholino represented as bar graph. Error bars represent SEM across four independent experiments.

**Fig S8:**
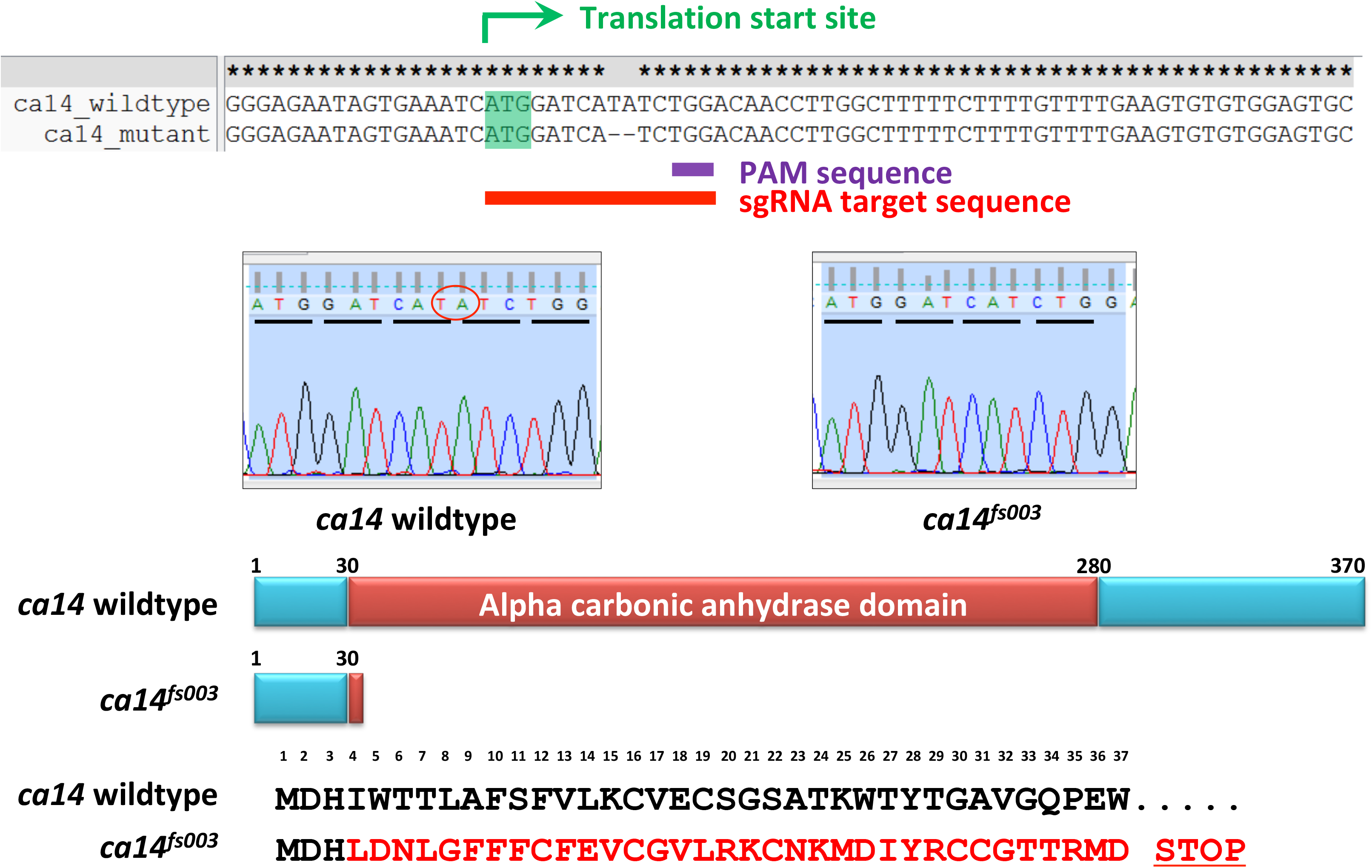
Schematic of zebrafish gene ca14 depicting that CRISPR target region and the observed mutation in ca14^fs003^.

### Supplementary Table 1

Excel sheet containing

A.Sequences of primers

B.qRT-PCR TAQMAN probes

C.siRNA and shRNA catalog numbers

D.Antibody catalog numbers and dilutions used in this study.

### Supplementary Information

**Fig S1:** 10% CO_2_ leads to extracellular acidification leading to decreased melanin content and increased proliferation

**Fig S2:** Gene expression data analysis from pigmentation models to identify putative effectors of melanocyte differentiation

**Fig S3:** Validation of Ca14 antibody used in the study

**Fig S4:** M-box sites in mouse and human Ca14 promoter and genic regions

**Fig S5:** mRNA expression of carbonic anhydrase 14 during pigmentation

**Fig S6:** Changes in CA14 correlate with pigmentation status of primary human melanocytes

**Fig S7:** ca14 morpholino causes pigmentation phenotype without affecting the survival of embryos

**Fig S8:** Schematic of zebrafish gene ca14 depicting the CRISPR target region and the observed mutation in ca14^fs003^.

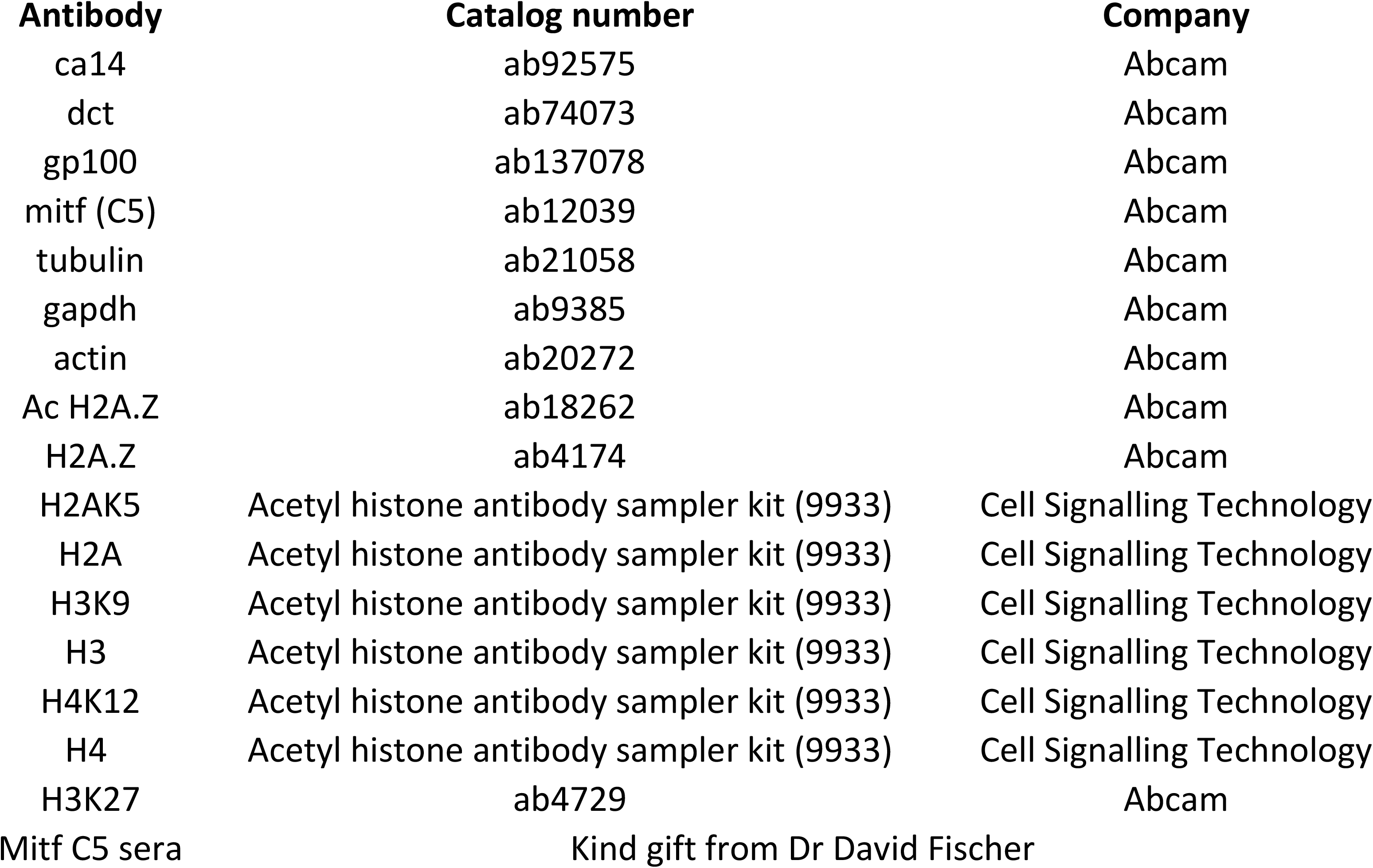
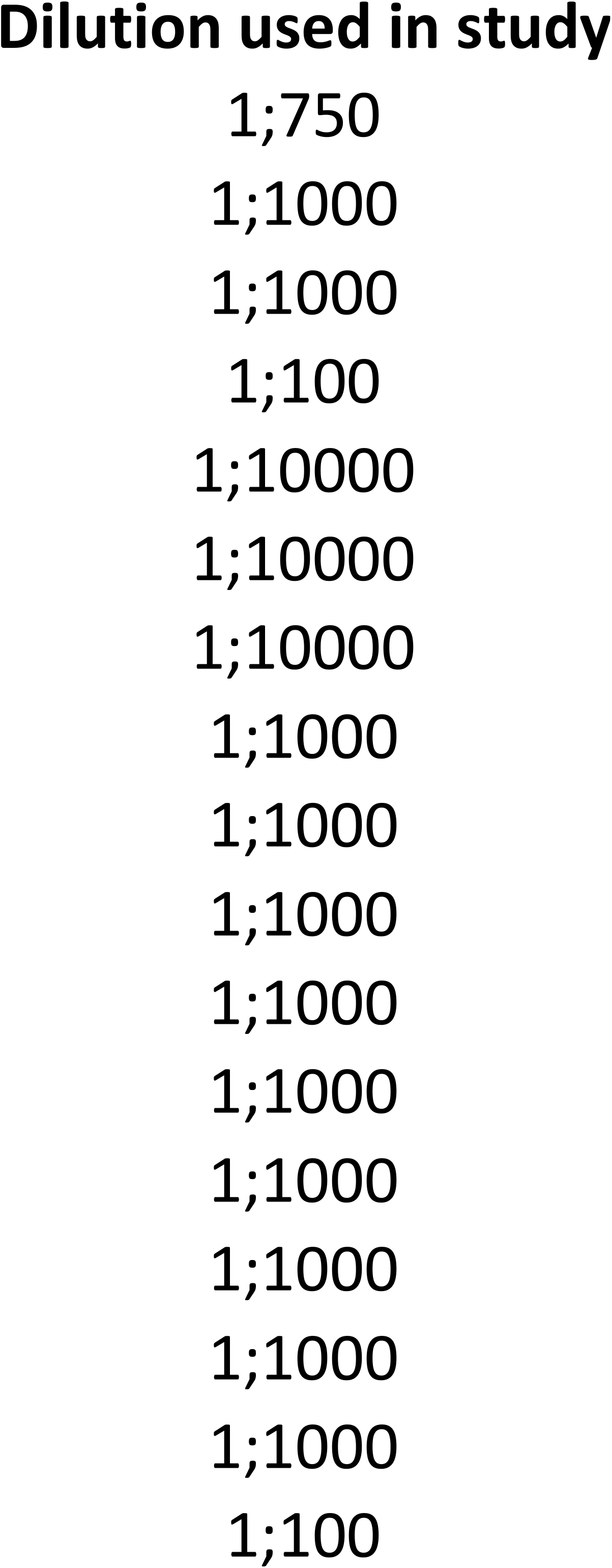

